# CD4^+^Foxp3E2^+^ regulatory T cell frequency predicts breast cancer prognosis and recurrence

**DOI:** 10.1101/2024.09.04.611142

**Authors:** Clorinda Fusco, Francesca Di Rella, Antonietta Liotti, Alessandra Colamatteo, Anne Lise Ferrara, Vincenzo Gigantino, Francesca Collina, Emanuela Esposito, Ivana Donzelli, Antonio Porcellini, Antonia Feola, Teresa Micillo, Francesco Perna, Federica Garziano, Giorgia Teresa Maniscalco, Gilda Varricchi, Maria Mottola, Bruno Zuccarelli, Bruna De Simone, Maurizio di Bonito, Giuseppe Matarese, Antonello Accurso, Martina Pontillo, Daniela Russo, Luigi Insabato, Alessandra Spaziano, Irene Cantone, Antonio Pezone, Veronica De Rosa

## Abstract

CD4^+^Foxp3^+^ regulatory T cells (Tregs) are key to maintain peripheral *self*-tolerance and suppress immune responses to tumors. Their accumulation in the tumor microenvironment (TME) correlates with poor clinical outcome in several human cancers, including breast cancer (BC). However, the properties of intratumoral Tregs remain largely unknown. Here, we found that a functionally distinct subpopulation of tumor-infiltrating Tregs, which express the Foxp3 splicing variant retaining exon 2 (Foxp3E2), is prominent in the TME and peripheral blood of hormone receptor- positive (HR^+^) BC subjects with poor prognosis. Notably, a comprehensive examination of the Tumor Cell Genome Atlas (TCGA) validated Foxp3E2 as an independent prognostic marker in all other BC subtypes. We found that FOXP3E2 expression underlies BCs with highly immune suppressive landscape, defective mismatch repair and a stem-like signature thus highlighting pathways involved in tumor immune evasion. Finally, we confirmed the higher immunosuppressive capacity of BC patients-derived Foxp3E2^+^ Tregs by functional assays. Our study suggests Foxp3E2^+^ Tregs might be used as an independent biomarker to predict BC prognosis and recurrence, and to develop super-targeted depletion-based immunotherapies.

**One-sentence summaries:** Foxp3E2^+^ Treg enrichment reflects an increased tumor-immune suppression and predicts prognosis and recurrence in breast cancer.

## Introduction

Immune surveillance against cancer is an important strategy for tracing, identifying, and eliminating growing tumor cells (*1–4*). Nonetheless, the immune system can shape tumor genomes by selecting neoantigen-depleted clones (i.e., immune editing) or promoting the accumulation of clones with an immune evasion strategy (i.e., immune escape), representing one of the main drivers of relapse (*5, 6*). Breast cancer (BC) is more resistant to immunotherapies than other solid tumors (*7, 8*), with a large window of recurrence spanning from months to decades after surgery (*9, 10*). Although the exact cause of this unusual recurrence pattern is still unknown, patients with luminal cancer typically have a better prognosis, whereas basal-like and HER2-enriched patients experience early relapses (within the first five years after diagnosis) (*11, 12*). However, the risk of late recurrence ranges from 10 to 41% in all BC subgroups, based on their primary tumor classification system (e.g. tumor-node-metastasis – TNM) (*13*), population-based data, and occasionally primary tumor gene expression profiles (*14*). Although the highest cumulative incidence has been observed among ER-positive patients, late recurrences also occur among those with ER-negative tumors (*15, 16*). It is, therefore, of paramount importance to identify novel prognostic biomarkers alongside with the causes of recurrence (*17*).

The interaction between tumor, stromal, and immune cells may promote metastatic progression and immune escape, challenging cancer immunotherapy efficacy (*5, 18*). Anti-tumor specific T cell responses arise in BC subjects but are halted by suppressive mechanisms established in the TME during tumor progression (*19*). CD4^+^CD25^+^ regulatory T cells (Tregs) expressing the Forkhead- box-p (Foxp)-3 transcription factor are enriched in the tumor microenvironment (TME) and associate with an invasive phenotype, reduced relapse-free and overall survival in several cancers (*20*), consistently with their role in suppressing effector cells. Transient depletion of Tregs *via* CD25, CTLA4, or CCR4 blockade results in improved clinical outcomes and increased anti-tumor specific immune responses (*19*). Foxp3^+^ Tregs variably infiltrate human BC and mainly correlate with reduced survival and poor prognosis (*1, 21–26*). Although their central function in tumor escape (*27*) and their role as therapeutic targets of immune checkpoint inhibitors (ICIs), the properties of intratumoral Tregs remain largely unknown and they are not a good prognostic marker for BC (*23, 28*). Published transcriptomic profiles indicate that tumor-infiltrating Tregs constitute a heterogeneous population (*29, 30*). Whether the tumor milieu imprints unique functional and transcriptional features to Tregs or whether distinct subsets of peripheral blood Tregs are differentially recruited within the tumor is still unclear (*31, 32*). Characterizing tumor-infiltrating Tregs will, therefore, be the key to find novel biomarkers and develop therapies that precisely target cells that block anti-tumor response without altering peripheral *self*-tolerance.

In humans, the master regulator of Treg development and function is *FOXP3* gene. It comprises 12 exons encoding multiple transcript variants, among which four are co-expressed at different levels in circulating Tregs, including the full-length (Foxp3FL) and those lacking exon 2 (Foxp3Δ2), which are generally more abundant (*33, 34*). Several reports uncover indispensable functions of the 105 base-pair region constituting *FOXP3* exon 2 (*FOXPE2*), highlighting a possible role of this region in regulating a transcriptional program that maintains Treg stability and immune homeostasis (*34–37*). In subjects with autoimmunity, we reported a selective reduction of Foxp3E2 splicing variants associated with impaired Treg suppressive function (*35*). Here, we study the distribution and function of Foxp3E2^+^ Tregs, both in the TME and peripheral blood of BC subjects in order to explore their connection with the molecular landscape of the primary tumor and patient prognosis.

## Results

### Elevated frequency of Foxp3E2+ Tregs in the tumor infiltrate and peripheral blood of HR+ BC subjects

High infiltration of Foxp3^+^ Tregs is expected to be associated with an unfavorable outcome in several cancers, but studies of breast cancer have led to highly discrepant findings (*38*). Here, we aimed at dissecting whether Tregs expressing different *FOXP3* variants could have a dominant role in breast cancer immune evasion (**Fig. 1a**). We analyzed the frequency of Foxp3^+^ (all FOXP3 transcript variants) and Foxp3E2^+^ Tregs (FOXP3 variants retaining exon2) in the peripheral blood (PB) and tumor-infiltrating lymphocytes (TILs) from two different cohorts of newly diagnosed, untreated ER^+^PR^+^(HR^+^)-HER2^-^ breast cancer (BC) and non-malignant breast fibroadenoma (BF) subjects (**Supplementary Table 1**). Freshly resected breast tissue was mechanically dissociated into a single-cell homogenate to enrich TILs (*39*). Flow cytometric analysis revealed a dominance of CD4^+^ T cells in BC, also confirmed by a lower CD8^+^/CD4^+^ ratio compared to BF tissue (0.86 *vs* 2.51) (**Fig. 1b**). In addition, BC tissue shows a more abundant infiltrate of Foxp3^+^ Tregs as compared to BF tissue and, a higher frequency of Treg cells is detected within the tissues as compared to peripheral blood (PB) from both BC and BF subjects (**Fig. 1c**). Importantly, we detected a significant enrichment of Foxp3E2^+^ Tregs in BC tissue (i.e. TIL-Foxp3E2^+^) compared to BF and PB (both from BC and BF subjects) (**Fig. 1d**). To estimate the relative frequency of Foxp3E2^+^ compared to the overall Treg compartment, we measured the Foxp3E2^+^/Foxp3^+^ ratio (herein defined E2 ratio) and found that Foxp3E2^+^ Tregs were more abundant both in the TME and PB of BC patients compared to BF (**Fig. 1e**). Furthermore, we compared the percentage and ratio of Foxp3E2^+^ and Foxp3^+^ Tregs in the TIL and PB of our BC cohort. The percentage of TIL- Foxp3E2^+^ Tregs was on average 8.33%, and the TIL-Foxp3^+^ represented 14.75% (**Fig. S1a**). The percentage of PB-derived Foxp3E2^+^ Tregs (PB-Foxp3E2^+^) and PB-Foxp3^+^ were, instead, 2.57% and 5.27% of the total CD4^+^ T cells, respectively (**Fig. S1a**). Notably the ratio between Foxp3E2^+^ and total Tregs in BC subjects was significantly higher in TIL compartment as compared to PB lymphocytes suggesting that Foxp3E2^+^ Tregs preferentially accumulate in the TME (mean E2 ratio is equal to 0.64 in TIL and 0.54 in PB) (**Fig. 1f**). Strikingly, the CD8^+^/Treg ratio was significantly lower in BC (both in PB and TIL) and inversely correlated with the percentage of TIL-Foxp3E2^+^ in the TME (*r* = -0.603, *P* = 0.001) (**Fig. 1g, h**), while no correlation was observed with the TIL- Foxp3^+^ (not shown). As the CD8^+^/Treg ratio is considered a reliable marker of anti-tumor specific T cell response (*34*), our data suggest that the Foxp3E2^+^ Treg subset mainly accounts for the suppression of the immune response to cancer. Immunohistochemical (IHC) staining and digital quantitative image analysis confirmed the higher infiltration of Foxp3^+^ and Foxp3E2^+^ Tregs in BC tissue compared to BF (**Fig. 1i-p**). Our results unveil for the first time a distinct prevalence of Foxp3E2^+^ Tregs in human BC that is not observed in non-malignant forms of breast tumors (i.e., BF) and inversely correlates with anti-tumor immune response.

**Figure 1.**
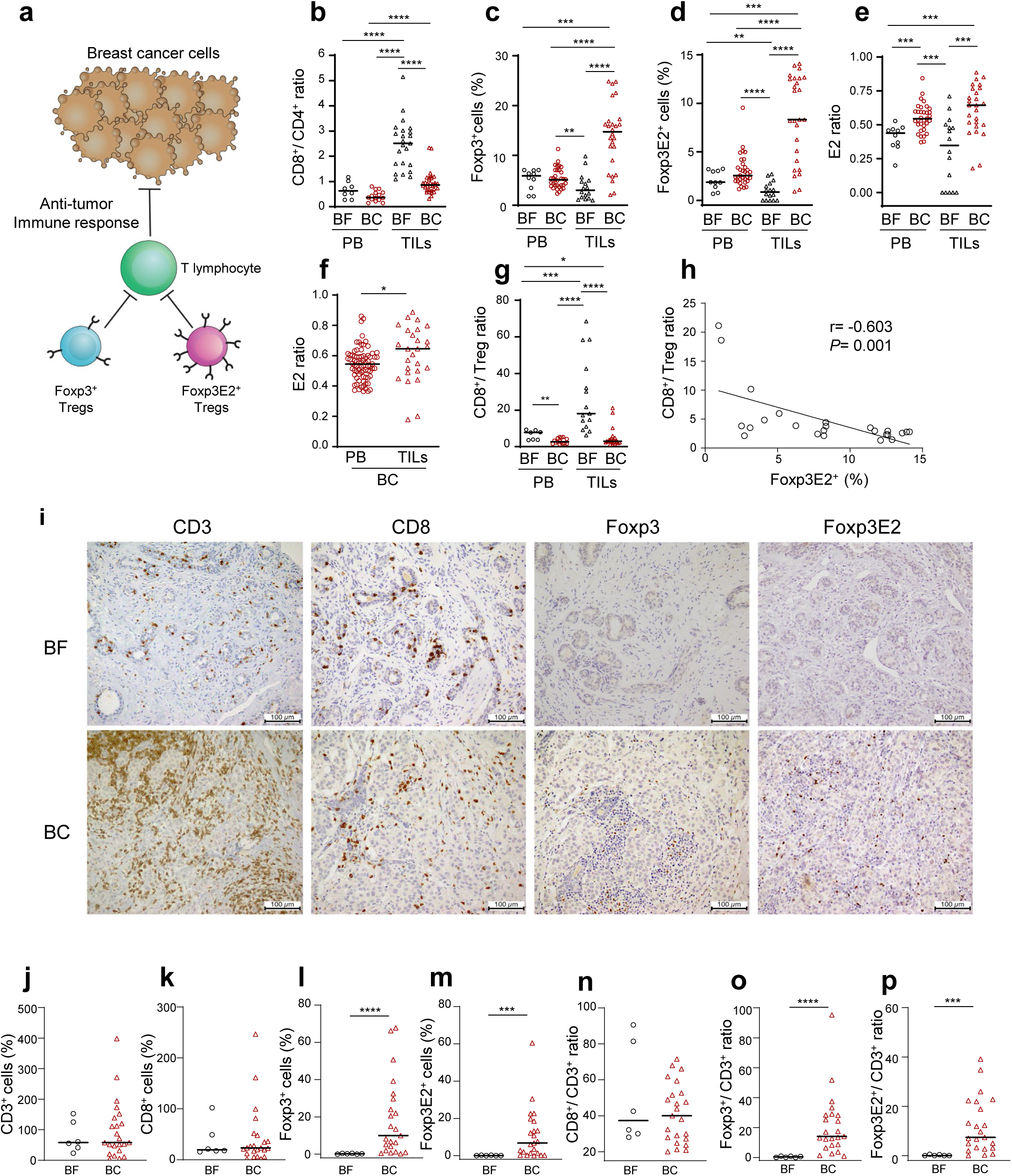
Characterization of the immune infiltrate in peripheral blood (PB) and primary tissue from breast cancer (BC) and fibroadenoma (BF) subjects. (a) Schematic representation of Foxp3^+^ and Foxp3E2^+^ Tregs in tumor immune escape. (b) CD8^+^/CD4^+^ ratio, % of (c) Foxp3^+^ and (d) Foxp3E2^+^ cells (gated on CD4^+^), (e-f) Foxp3E2^+^/Foxp3^+^ ratio (E2 ratio) and (g) CD8^+^/Treg ratio in peripheral blood (PB – represented as dots) and tumor-infiltrating lymphocytes (TILs – represented as triangles) from BF (white empty) and BC (red empty) subjects. In panel (b-g), represented data are for BF at least *n* = 7 and *n* = 15 and for BC at least *n* = 15 and *n* = 24 (respectively for PB and TILs), *n* = 77 for PB and *n* = 26 for TILs. (h) Correlation between % of Foxp3E2^+^ and CD8^+^/Treg ratio in TILs from BC subjects (*n* = 24). (i) Representative immunohistochemical staining of primary BC and BF tissue showing CD3^+^, CD8^+^, Foxp3^+^ and Foxp3E2^+^ cells. Immunohistochemistry-based quantification of (j) % of CD3^+^, (k) CD8^+^, (l) Foxp3^+^, (m) Foxp3E2^+^ cells, (n) CD8^+^/CD3^+^ ratio, (o) Foxp3^+^/CD3^+^ ratio and (p) Foxp3E2^+^/CD3^+^ ratio [respectively white dots (*n* = 6) for BF and red triangles (*n* = 23) for BC subjects]. Data are presented as Median values. Each data point represents a different individual (i.e., independent biological samples) (a-e, g, h, j-p) or experimental replicates (f). Statistical analyses were performed by using Mann-Whitney *U*-test (two tails) (a-g, j-p) and Spearman r correlation test (h). * *P*≤ 0.05; \*\**P*≤ 0.01; \*\*\**P*≤ 0.005; \*\*\*\**P*≤ 0.0001.

### *FOXP3E2* transcript levels in BC tissue mark an immunosuppressive landscape and correlate with reduced overall survival

To determine whether the increased percentage of Foxp3E2^+^ Tregs is associated with BC prognosis in general, we examined RNAseq data from about one thousand subjects (990 breast cancer tissues and 112 tumor-adjacent normal tissues) in the TCGA Splicing Variant Database (TSVdb) that includes information on alternative splicing (*40*). We found that primary breast cancers (69.2% HR^+^HER2^-^, 12.6% HR^+^HER2^+^, 18.2% HR^-^HER2^-^) expressed higher levels of *FOXP3* transcripts compared to normal breast tissue (NT) (64.00 *vs* 13.40) (**Fig. 2a**). However, *FOXP3* transcript levels did not correlate with patient overall survival when BC subjects were stratified either on their median value (Q2) or on their upper quartile range (Q3) (**Fig. 2b, c, Fig**. **S1b, c**). Thus, we measured the expression of the 5 different *FOXP3* isoforms (schematically represented in **Fig. S1d** and reported in the UCSC bank (*33*)), and we found that 4 of them were upregulated in BC tissue compared to NT (**Fig. S2a-d**) but none correlated with overall survival (**Fig. S2e-l**). Then, we estimated the ratio of *FOXP3* exon 2-containing transcripts relative to the other variants and stratified BC subjects into low- (< Q3 = 0.09) and high- (> Q3 = 0.29) E2 ratio groups (**Fig. 2d, e**). This analysis clearly shows that the difference in *FOXP3E2* expression between the two groups was inversely correlated with patient overall survival (log-rank *P* = 0.01, Chisq = 6.2) (**Fig. 2f, g**). No difference in total *FOXP3* expression between the two groups was observed (**Fig. S3a**); also, we did not find correlation when BC subjects were stratified on the median value of the E2 ratio (**Fig. S3b-d**). Notably, TCGA also included HER2^+^ as well as the most aggressive triple negative tumors, thus suggesting a general association between the enrichment of Foxp3E2^+^ Tregs within the tumor and breast cancer prognosis.

**Figure 2.**
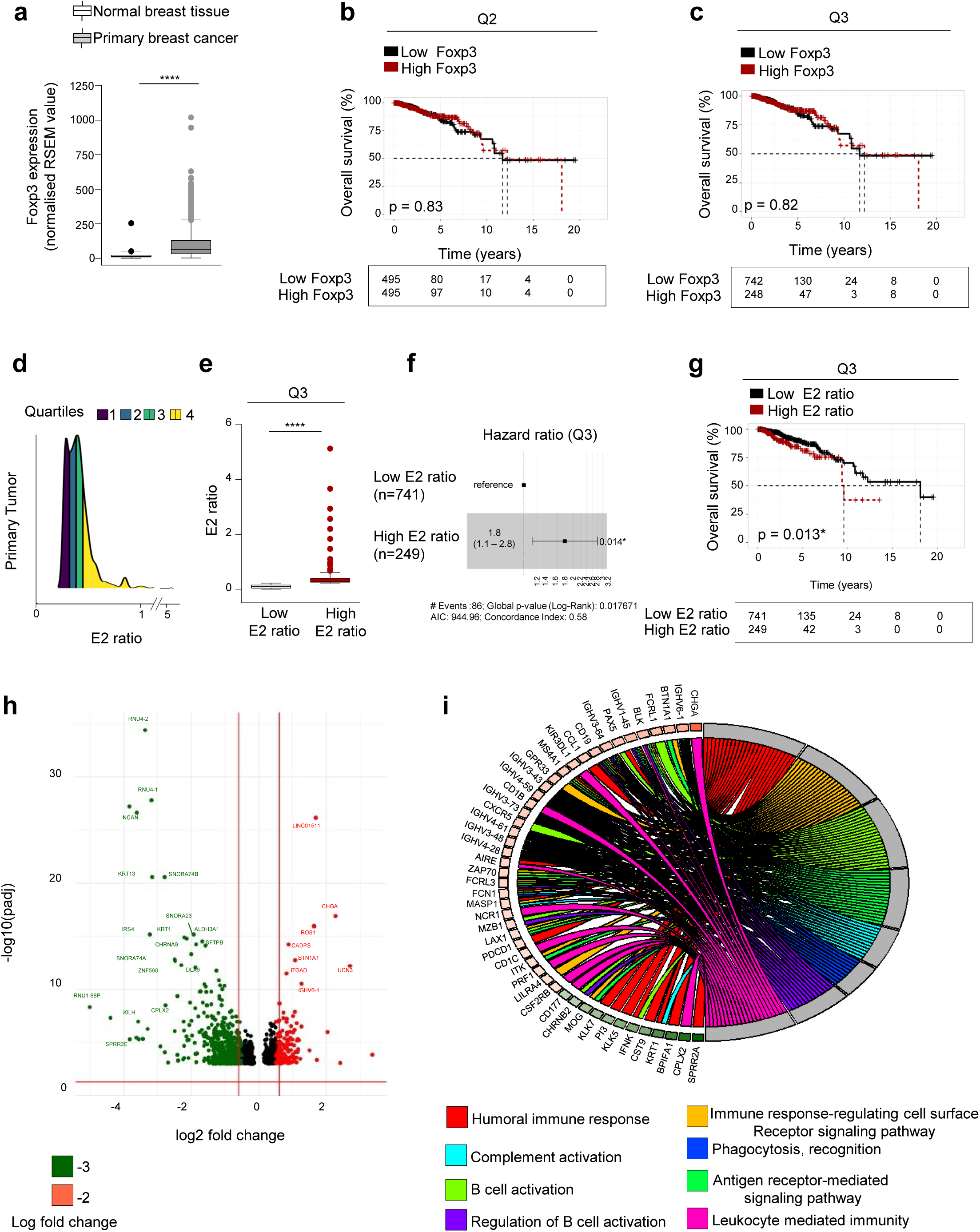
Foxp3E2 transcript analysis from primary breast cancer (BC) tissues delineates a subgroup of subjects with poor prognosis and a distinct gene expression profile. **(a)** Foxp3 transcripts in normal (*n* = 112) and primary breast cancer (*n* = 990) tissues. Data represent normalized RSEM value obtained by RNAseq analysis of datasets in the TCGA Splicing Variant Database. **(b, c)** Kaplan-Meier survival curve of BC subjects stratified into low- and high-Foxp3 expression levels within the primary tumor based on its Q2 (*n* = 495 and 495) or Q3 (*n* = 742 and 248) value. **(d)** Interquartile distribution of the Foxp3E2^+^/Foxp3^+^ ratio calculated in the primary BC tissue (*n* = 990). **(e)** BC subjects were stratified into low- (*n* = 741) and high- (*n* = 249) Foxp3E2^+^/Foxp3^+^ ratio (E2 ratio) according to the Q3 value cut-off. **(f)** Hazard Ratio (HR = 1.8, CI 1.1 – 2.8, Cox *P* = 0.014) and **(g)** Kaplan-Meier survival curve of low- (*n* = 741) and high- (*n* = 249) E2 ratio BC subjects according to Q3 value cut-off. **(h)** Volcano plot of Differentially Expressed Genes (DEGs) obtained by applying a threshold of log2 foldchange > ± 0.05 (x-axis) and a p-adj < 0.001 (y-axis) in the two groups of BC subjects. Dots represented single genes: 179 upregulated (red), and 523 downregulated (green) in the high-ratio BC group. **(i)** Circular composition overview plot for selected gene ontology pathways (represented in different colors) overrepresented among DEGs in high- *vs* low-ratio BC groups. Gene Ontology (GO) analysis was performed by DAVID (Database for Annotation, Visualization and Integrated Discovery) database Gene color scale indicates the relevant fold change values (red -upregulated, green - downregulated). Data are presented as Median values. Statistical analyses were performed by using Mann-Whitney *U*-test (two tails) **(a, e)**, Multivariate Cox regression model reference **(b, c, g)**, and log-rank test **(f)**. \*\*\*\**P*≤ 0.0001.

To gain insights into the nature of the local TME (*41, 42*), we characterized gene expression patterns of high- and low- Foxp3E2^+^/Foxp3^+^ ratio BC groups. Analysis of differentially expressed genes (DEGs) identified 702 DEGs (523 downregulated and 179 upregulated genes) (**Fig. 2h**). Gene ontology revealed a significant enrichment of genes belonging to immunoregulatory pathways in the BC group showing high Foxp3E2^+^/Foxp3^+^ ratio. These immunoregulatory genes included humoral immune response, complement activation and antigen receptor-mediated signaling (**Fig. 2i**). Among all, the upregulation of *BTN1A1*, *FCRL1*, *CXCR5*, *AIRE*, *ZAP70* was noteworthy (**Fig. 2i**) indicating a dominant immunosuppressive signature (*43, 44*). Consistently, GSEA-KEGG analysis showed 5 sub-gene sets activated in the high E2 ratio BC group (chemokine signalling, hematopoietic cell lineage, glycerolipid metabolism, cAMP signaling and neuroactive ligand-receptor interaction (*45*)) (**Fig. S3e**). Upregulation of *WNT3a*, *ESRG*, *NANOGP1*, *NEFL*, instead, suggested the acquisition of stem cell-like features (*46–49*) (**Fig. S4**). Overall, these analyses reveal that Foxp3E2 marks a distinctive group of BC subjects characterized by worst clinical outcomes (i.e., lower survival) and likely associated with increased immunosuppression and stemness.

### BC tumors with high Foxp3E2^+^/Foxp3^+^ ratio show greater heterogeneity

To better understand the relationship between high E2 ratio and poor BC clinical outcome, we investigated mutations in cancer driver genes. To this aim, we first characterized genome variants in the high- and low-E2 ratio BC groups. We observed comparable tumor mutational burden in the two groups, with *PIK3CA* and *TP53* mutations dominating the landscape (**Fig. 3a, b, Fig**. **S5a, b**) consistently with previous reports (*50*). Other genes, however, harbored coding mutations in at least 6% of the samples: *TTN*, *MUC16*, *MAP3K1*, *KMT2C*, *GATA3*, *SYNE1* and *FLG* (**Fig. 3a, b**). Well-known germline mutations in *BRCA1* and *BRCA2* were identified in less than 5% of BC subjects of both groups (**Fig. S6**). We then examined pairwise associations between somatic events to explore co-mutations and mutual exclusivity patterns and found mutual exclusivity between *TP53*, *GATA3* and *CDH1* mutations in both BC patient groups, suggesting they might have originated from similar ancestral clones. Interestingly, the PI3K/Akt co-mutation marked specifically the high Foxp3E2^+^/Foxp3^+^ ratio BC group (false discovery rate (FDR) <0.05) (**Fig. 3c, d**), further suggesting cancer stem-like features (*51, 52*). Furthermore, we detected a lower frequency of co-mutations in the high Foxp3E2^+^/Foxp3^+^ ratio BC group despite the comparable tumor mutation burden (**Fig. 3c, d**). The latter might reflect a sub-clonal heterogeneity that has been associated with therapy resistance and tumor shaping (*53–56*). Overall, this suggests that Foxp3E2^+^ Tregs mark a subgroup of tumors with greater intratumor heterogeneity.

**Figure 3.**
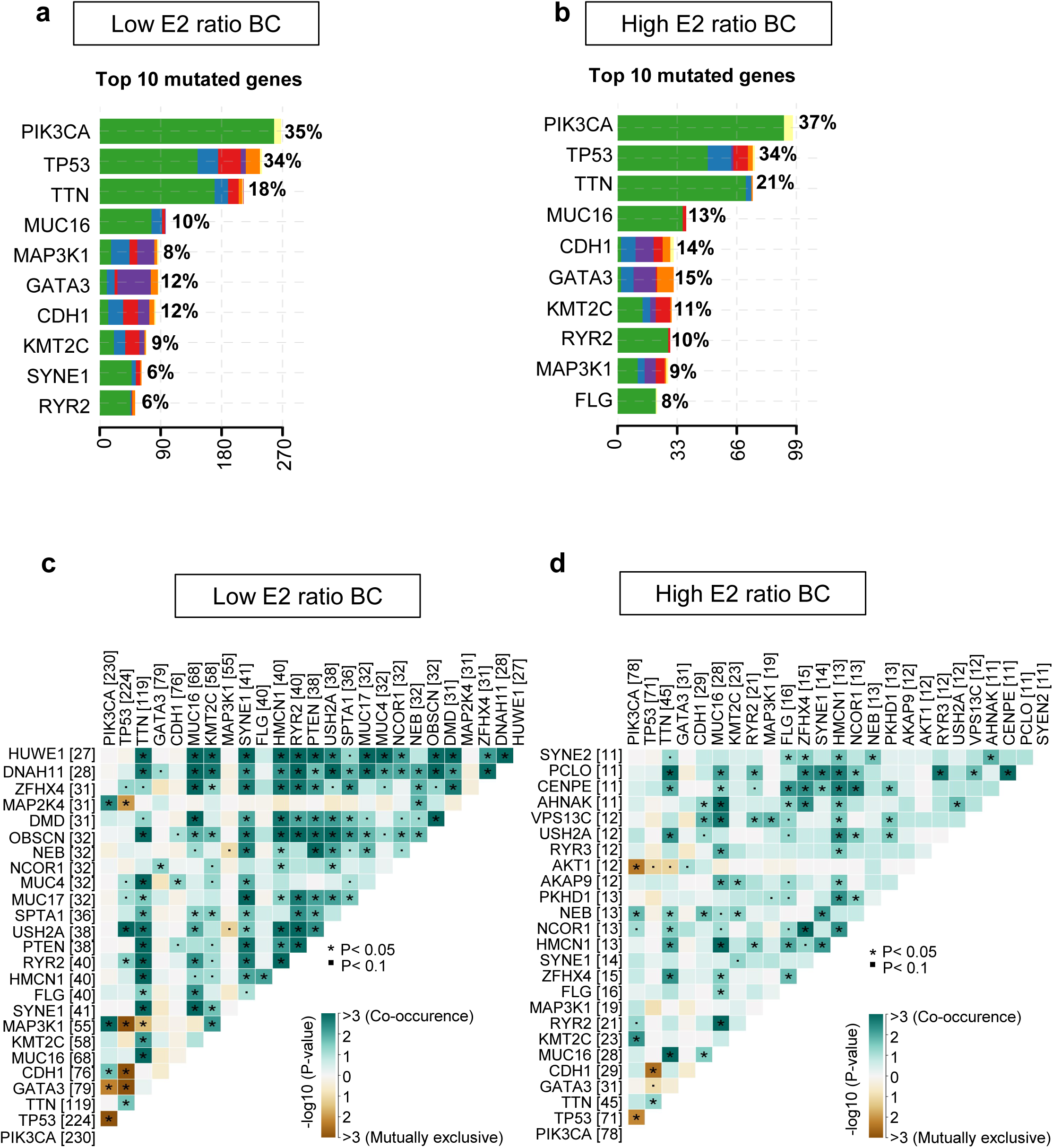
High intra-tumor heterogeneity characterizes the BC group with high Foxp3E2^+^/Foxp3^+^ ratio. **(a, b)** Summary of top 10 mutated genes in BC subjects with **(a)** low- and **(b)** high- E2 ratio. **(c, d)** Somatic interaction analysis between gene pairs showing co-occurring mutations (green squares) and mutually exclusive mutations (brown squares). The intensity of the color is proportionate to the – log10 (*P*-value). Statistical analyses were performed by using Fisher’s exact test.

### BC tumors with high Foxp3E2^+^/Foxp3^+^ ratio show mutational signatures associated with defective DNA mismatch repair and strong immunosuppressive response

The sub-clonal nature of the high E2 ratio BC group suggests a tumor evolution and selection (*57, 58*), which implies different molecular mechanisms including spontaneous and enzymatic deamination of the cytosine base (*59, 60*). To gain insights into the dynamics of the mutational signature that shapes both BC groups, we interrogated COSMIC mutational signatures that have been associated with specific pathways (*30*). Signatures associated with spontaneous deamination of 5-methylcytosine and APOBEC cytidine deaminase were detected in both the high- and low-E2 ratio BC groups (**Fig. 4a, b**). Enrichment analysis of APOBEC motif (i.e. tCw motif primarily associate with C>T transitions driven by APOBEC cytidine deaminase activity) in the high and low Foxp3E2^+^/Foxp3^+^ ratio BC groups showed a similar prevalence of tCw mutations (34% APOBEC *vs* 9% non-APOBEC in the low-E2 ratio BC group; 38% APOBEC *vs* 9% non-APOBEC in the high-E2 ratio BC group) (**Fig. 4c, d**), with no changes in the global DNA methylation (**Fig. S7a**). Defects in polymerase POLE, which occur in ultra-hypermutators, have instead been observed only in the low-E2 ratio BC group (**Fig. 4e**), thus confirming the high tumor mutational burden associated with the increased presence of immunogenic neoantigens (*54*).

**Figure 4.**
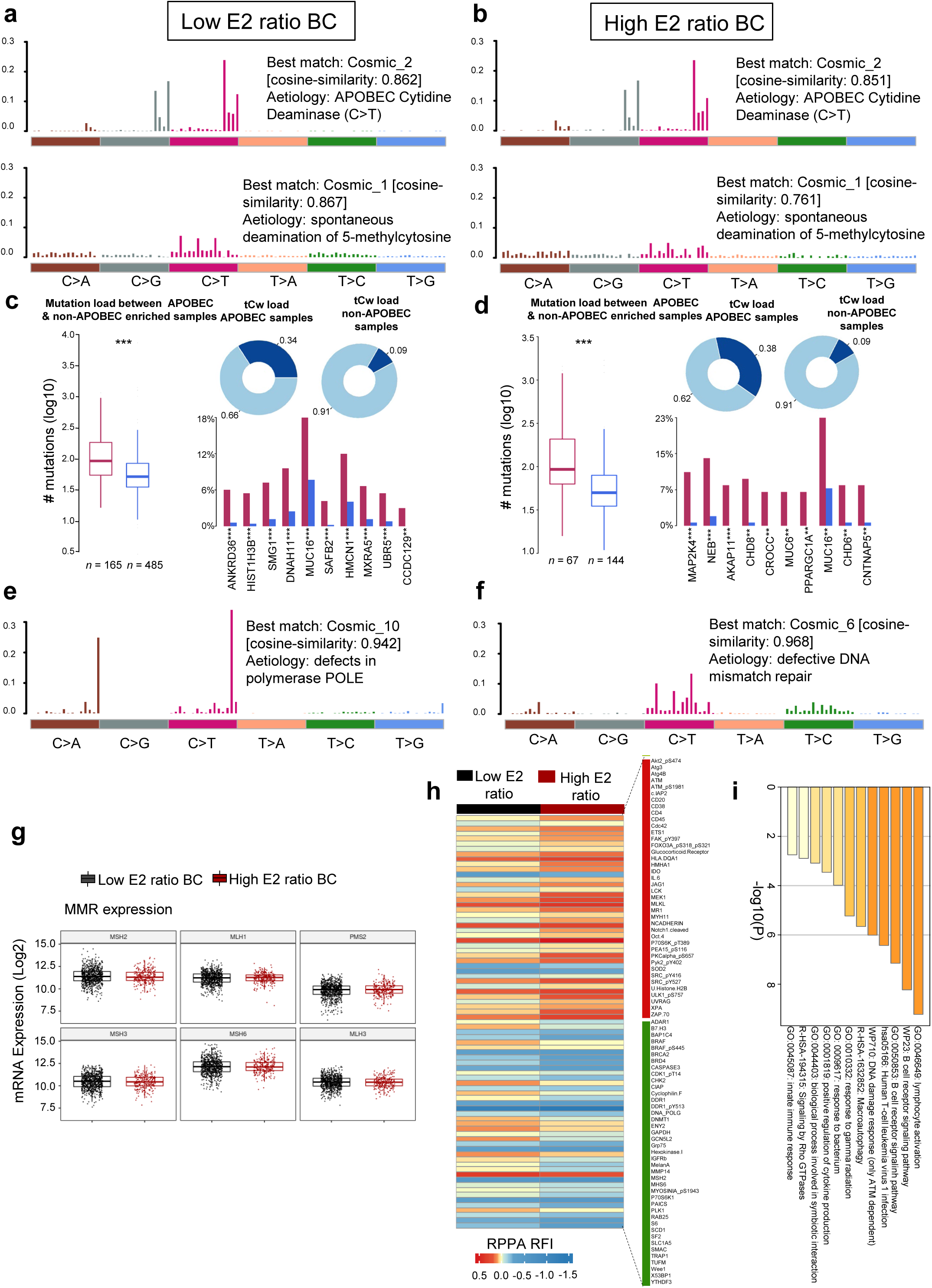
Defective DNA mismatch repair and specific mutational signatures in BC subjects with high Foxp3E2^+^/Foxp3^+^ ratio. Mutational signatures identified in BC subjects with **(a, e)** low- and **(b, f)** high- Foxp3E2^+^/Foxp3^+^ ratio, respectively. The y-axis indicates exposure of 96 trinucleotide motifs to overall signature. In each plot, we report the best match against validated COSMIC signatures and cosine similarity value alongside the proposed etiology. **(c, d)** APOBEC enrichment analysis in BC subjects with **(c)** low- and **(d)** high-E2 ratio. Box plots (left panels) show differences in mutation load between APOBEC-enriched and nonenriched samples. Donut plots (upper panels) display the proportion of mutations in tCw context. Bar plots (lower panels) show the top 10 differentially mutated genes between APOBEC-enriched and non-APOBEC-enriched samples. **(g)** Box plots reporting the expression profiles of MMR-relative genes (MLH1, MLH3, MSH2, MSH3, MSH6, PMS2). **(h)** Supervised hierarchical clustering analysis of TCGA-BC RPPA results using an ANOVA FDR p-value threshold lower than 0.05. Based on this threshold, 81 probes were differentially altered in the high-E2 ratio group, with 40 probes upregulated (red bar) and 41 downregulated (green bar). **(i)** Enrichment analysis of the differently expressed probes using Metascape. Statistical analyses were performed by using the Wilcoxon rank-sum test and Fisher’s exact test **(a, b, e, f)**; \*\*\**P*≤ 0.005.

Strikingly, the signature associated with defective DNA mismatch repair (dMMR) was specific for the high-E2 ratio BC group (**Fig. 4f**). MMR is a fundamental DNA repair pathway essential to maintain genome stability during cellular replication (*61*) and defects have been considered driver of endocrine treatment resistance in 15-17% of ER^+^/HER2^−^ BC subjects (*62, 63*). To better dissect this pathway, we evaluated gene and protein expression of MMR-associated factors. We did not observe changes in MMR gene expression (e.g. *MLH1*, *MLH3*, *MSH2*, *MSH3*, *MSH6*, *PMS2*) (**Fig. 4g**). However, analysis of reverse phase protein array (RPPA) showed, instead, high levels of ATM, ATM_pS1981, UVRAG, XPA and low levels of BRCA2, CHK2, DDR1, DDR1_pY513, DNA PolG, MSH2, MSH6, Wee1, X53BP1 and DNMT1 in the high Foxp3E2^+^/Foxp3^+^ BC subjects (**Fig. 4h**), suggesting that DNA Damage Response (DDR) might be specifically dysregulated in these patients.

Notably, BC patients with high Foxp3E2^+^/Foxp3^+^ ratio also showed higher expression of immuno-modulatory signatures as compared with the low-E2 ratio group (i.e., *CD20*, *CD38*, *CD4*, *CD45*, *IL6*, *JAG1*, *ZAP70*) (**Fig. 4h**). This increase in immunomodulatory pathways was also confirmed by enrichment analysis of the differentially expressed probes using Metascape (*64*) (**Fig. 4i**). Notably, immune cell deconvolution shows only a slight increase in the number of endothelial cells between the two subgroups with no difference in the immune cell compartments (**Fig. S7b**). Altogether our analyses unveil that Foxp3E2 marks a subgroup of breast tumors characterized by defective DNA damage repair and strong immunosuppressive signature.

### Increased immune checkpoint expression and stronger suppressive capability is associated with Foxp3E2 expression in Tregs of HR^+^ BC

As both our *ex-vivo* analyses of HR^+^ BC subjects and TCGA data mining suggest an association between Foxp3E2^+^ Tregs and immunosuppression in the TME, we evaluated their immunosuppressive function. To this aim, we initially checked the expression of a range of co- inhibitory molecules – known to modulate tumor immune responses and upregulated in tumor- infiltrating Tregs (*15, 21, 22, 65*) (e.g., immune checkpoints (ICs), such as CTLA-4, PD-1 and TIGIT) – in TIL and PB-derived CD4^+^ T cells from BC patients (**Fig. 5**, **Fig. S8**). We found that TIL-Foxp3E2^+^ Tregs have increased percentage of Helios and CCR8 and higher levels of Helios and CTLA-4 than TIL-Foxp3^+^ Tregs. Moreover, when compared to PB, TIL-Foxp3E2^+^ Tregs show increased expression of Helios, ICOS, CTLA-4, PD-1, TIGIT and CCR8, and higher proliferative capacity, as pointed out by Ki67^+^ frequency (**Fig. 5**, **Fig. S8**). Interestingly, co-expression of CTLA- 4 and PD-1 or TIGIT and CCR8 was higher in TIL-Foxp3E2^+^ than in PB-Foxp3E2^+^ Tregs (**Fig. 5**), thus suggesting that this Treg subpopulation exerts a dominant role in cancer evasion/immunosuppression. This is particularly relevant as elevated CCR8 expression in TIL- Tregs are related to poor prognosis in several cancer types (*19, 66*). Moreover, the increased expression of Helios and ICOS revealed that Foxp3E2^+^ Tregs infiltrating the tumor had a hyper- activated phenotype. Also, the evidence that pS6 levels – reflecting mTOR kinase activity – were reduced specifically in TIL-Foxp3E2^+^ compared to the PB-counterpart, suggests a detrimental role for the mTOR pathway in the suppression of anti-tumor response. Overall, the increased expression of Helios and IC in the Foxp3E2^+^ Tregs proposes that they might have a higher immunosuppressive capacity in cancer. We, indeed, tested this hypothesis through an *in vitro* CFSE-based suppression assay by culturing blood-derived conventional T cells (Tconvs) with autologous Tregs from BC subjects. Tregs from BC patients show stronger suppressive capacity compared to Tregs from age- matched healthy female donors (HD) (**Fig. 6a**). When compared to HD, BC subjects displayed increased E2 PB-ratio (0.54 *vs* 0.49) (**Fig. 6b**), and higher frequency of TIGIT^+^, CCR8^+^, TIGIT/CCR8 and CTLA-4/PD-1 double positive PB-Foxp3E2^+^ Tregs (**Fig. 6c**, **Fig. S9**). Interestingly, the lower expression of ICOS in PB-Foxp3E2^+^ Tregs of BC subjects suggested a preferential recruitment of ICOS^+^Foxp3E2^+^ Tregs in the TME, as ICOS levels were higher in TIL- Foxp3E2^+^ Tregs compared to the peripheral blood (**Fig. 5**, **Fig. 6c**). Notably, the median of the E2 PB-ratio within the BC cohort (Q2 = 0.545) represents a “hub value” almost coincident with that corresponding to the Q3 value of HD (Q3 = 0.546) (**Fig. 6d**), suggesting E2 PB-ratio might as well be associated with stronger immune suppression.

**Figure 5.**
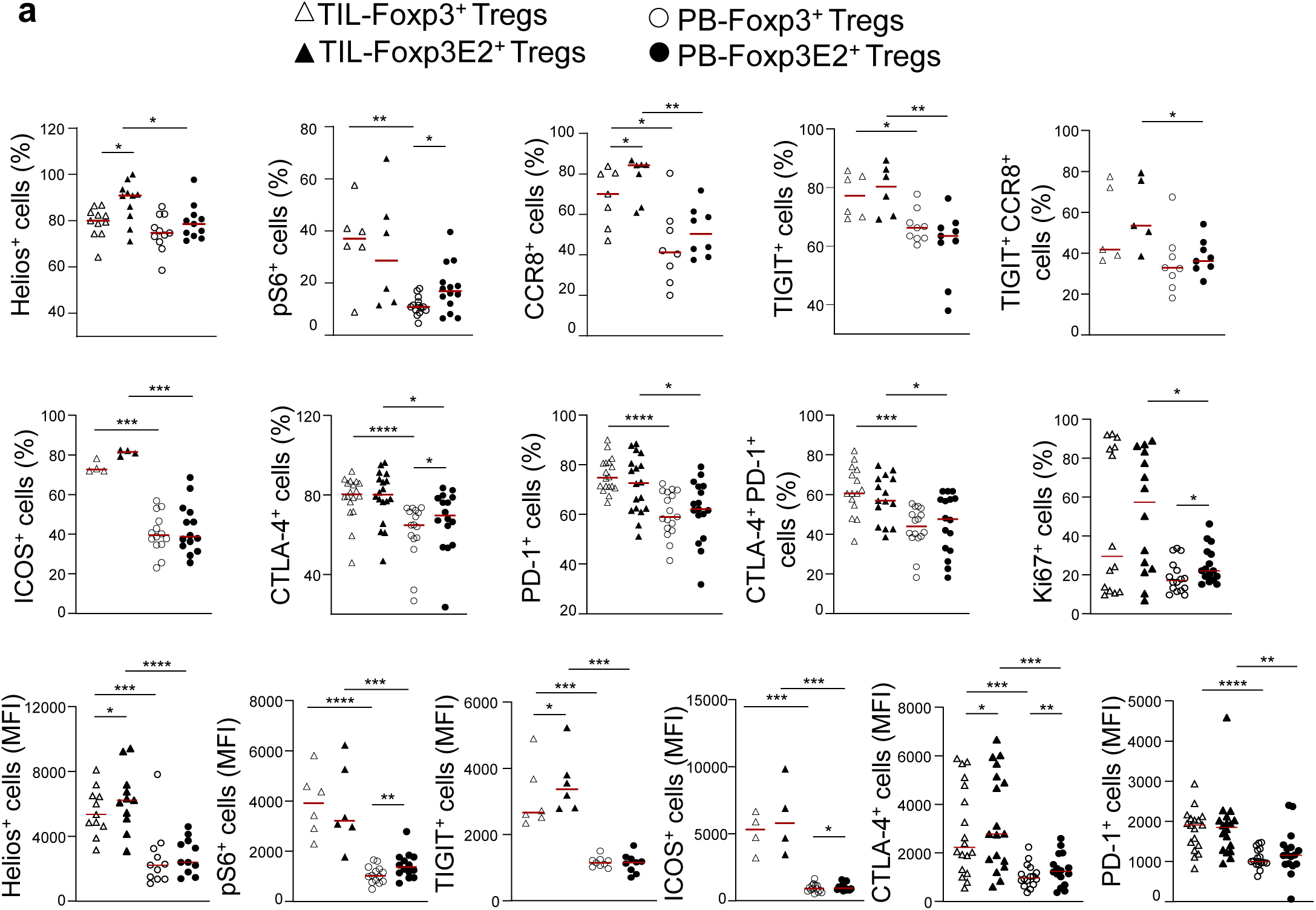
Highly immunosuppressive Foxp3E2^+^ Tregs preferentially accumulate in TILs of newly diagnosed HR^+^ BC subjects. Cumulative data of flow cytometry analysis showing cell percentage and mean fluorescence intensity (MFI) of Helios^+^, pS6^+,^ CCR8^+^, TIGIT^+^, ICOS^+^, CTLA-4^+^, PD-1^+^ and Ki67^+^ cells (gated on CD4^+^Foxp3^+^ and CD4^+^Foxp3E2^+^) in freshly isolated TILs (at least *n* = 4) and PB (at least *n* = 9) from BC subjects. Data are presented as Median values. Statistical analysis was performed by using Wilcoxon and Mann-Whitney *U*-test (two tails); \**P*≤ 0.05; \*\**P*≤ 0.01; \*\*\**P*≤ 0.005; \*\*\*\**P*≤ 0.0001.

**Figure 6.**
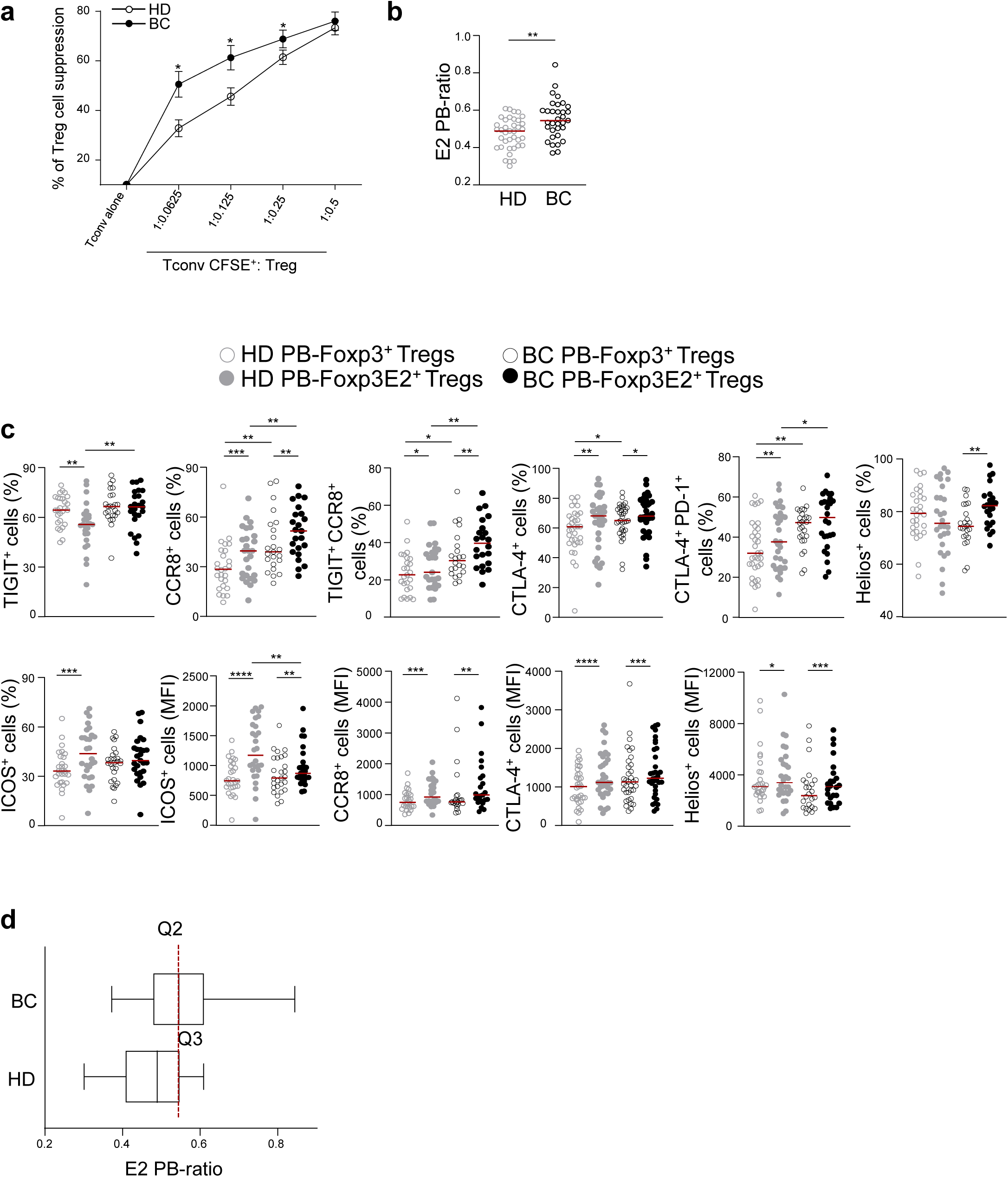
Higher immune checkpoint expression in Tregs from HR^+^ BC subjects correlates with increased Foxp3E2^+^/Foxp3^+^ ratio and peripheral Treg suppressive function. **(a)** Percentage of suppression of Tregs in co-culture with CFSE-labeled Tconvs at different ratios, from HD (*n* = 20) and BC (*n* = 16) subjects. **(b)** Cumulative data calculated by flow cytometry quantification of the E2 ratio (evaluated on CD4^+^Foxp3^+^ and CD4^+^Foxp3E2^+^ Tregs) from PB of HD (*n* = 38) and BC (*n* = 33) subjects. **(c)** Percentage of TIGIT^+^, CCR8^+^, TIGIT^+^/CCR8^+^, CTLA- 4^+^, CTLA-4^+^/PD-1^+^, Helios^+^ and ICOS^+^ Tregs and MFI of ICOS, CCR8, CTLA-4 and Helios on CD4^+^Foxp3^+^ and CD4^+^Foxp3E2^+^ Tregs from freshly isolated PB of HD (at least *n* = 25) and BC (at least *n* = 22) subjects. **(d)** E2 PB-ratio (median, minimum to maximum values, and quartiles) from BC (*n* = 33) and HD (*n* = 38) subjects. Each symbol shows independent biological samples **(b-d)** or experimental replicates **(a)**. Data are presented as Median values. Statistical analysis was performed by using Wilcoxon and Mann-Whitney *U*-test (two tails); \**P*≤ 0.05; \*\**P*≤ 0.01; \*\*\**P*≤ 0.005; \*\*\*\**P*≤ 0.0001.

Then we observed that Treg peripheral suppression in BC subjects directly correlated with the E2 TIL-ratio (*r* = 0.66, *P* = 0.021) (**Fig. 7a**). Specifically, BC subjects with high E2 TIL-ratio (≥ 0.64) showed higher peripheral Treg suppression than the ones with a low ratio (< 0.64) (**Fig. 7b**). The more TIL-Foxp3E2^+^ Tregs they had, the greater was their peripheral suppressive capacity. Importantly, we observed that the E2 PB-ratio strictly mirrored the E2 TIL-ratio, as high E2 TIL- ratio BC subjects also exhibited higher E2 PB-ratio (**Fig. 7c**). Consistently, Tregs from BC patients with higher E2 PB-ratio showed increased suppressive activity compared to the low E2 PB-ratio group (**Fig. 7d**). Furthermore, Foxp3E2^+^ Tregs from the high E2 PB-ratio BC group displayed an immune phenotype distinct from the Foxp3^+^ Tregs, with enhanced expression of Helios, ICOS, CTLA-4, CCR8, and co-expression of CTLA-4/PD-1 and TIGIT/CCR8 (**Fig. 7e**, **Fig. S10a**) thus mirroring the hyperactivated phenotype observed in TIL-Foxp3E2^+^ Tregs (**Fig. 5**). Strikingly, Foxp3E2^+^ Tregs from the high E2 PB-ratio expressed low pS6 levels compared to Foxp3E2^+^ Tregs from the low E2 PB-ratio group (**Fig. 7e**), according to what previously observed in TIL-Foxp3E2^+^ *versus* TIL-Foxp3^+^ (**Fig. 5**). This could further support that mTOR activity perturbs the suppression of antitumor-specific immune response.

**Figure 7.**
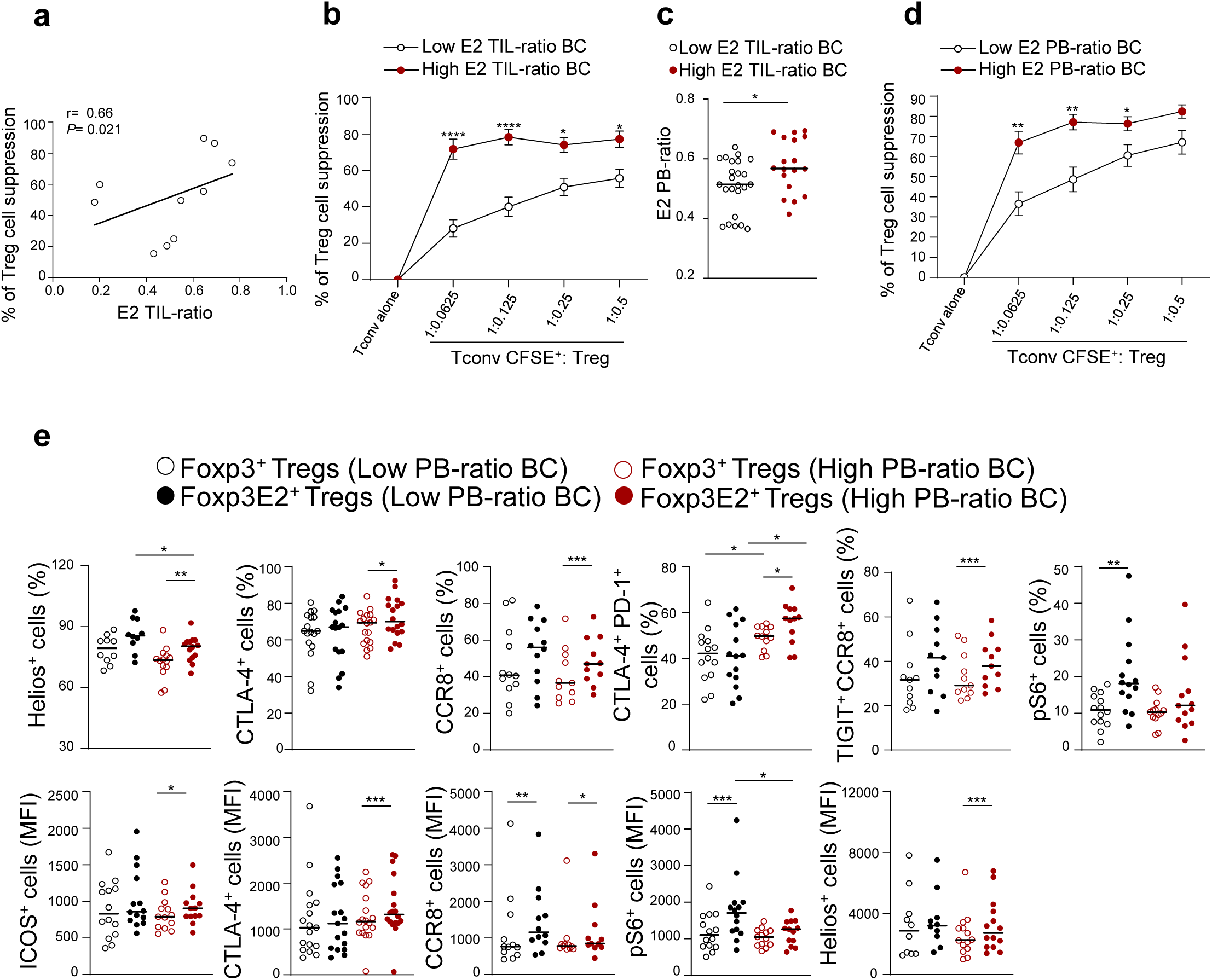
Increased peripheral Treg suppressive function and immune checkpoint expression in Tregs from BC subjects with higher Foxp3E2^+^/Foxp3^+^ TIL- and PB-ratio. **(a)** Correlation between the E2 TIL-ratio and the percentage of peripheral suppression of Tregs from BC subjects (*n* = 10). **(b)** Percentage of suppression of Tregs from BC subjects with high (*n* = 6) and low (*n* = 6) E2 TIL-ratio BC subjects at different proportions of Treg/Tconv cells. **(c)** E2 PB-ratio from BC subjects with high (*n* = 18) and low (*n* = 24) TIL-ratio. **(d)** Percentage of Treg suppression in BC subjects divided into high- (*n* = 7) and low- (*n* = 10) Foxp3E2^+^/Foxp3^+^ PB-ratio. **(e)** Cumulative data calculated by flow cytometry quantification showing the percentage of Helios^+^, CTLA-4^+^, CTLA-4^+^PD-1^+^, CCR8^+^, TIGIT^+^CCR8^+^ and pS6^+^ cells and MFI (Helios, CTLA-4, CCR8, pS6 and ICOS) gated on CD4^+^Foxp3^+^ and CD4^+^Foxp3E2^+^ Tregs from peripheral blood of BC subjects with high- (at least *n* = 11) and low- (at least *n* = 13) E2 PB-ratio BC subjects. Each symbol shows independent biological samples **(a-c, e)** or experimental replicates **(b, d)**. Data are presented as Median values. Statistical analysis was performed by using Wilcoxon and Mann-Whitney *U*-test (two tails); \**P*≤ 0.05; \*\**P*≤ 0.01; \*\*\**P*≤ 0.005; \*\*\*\**P*≤ 0.0001.

Overall, our data indicate that the E2 ratio in the peripheral blood reflects the infiltration of highly immunosuppressive Foxp3E2^+^ Tregs in the TME.

### Foxp3E2^+^/Foxp3^+^ ratio predicts the prognosis in newly-diagnosed HR^+^ BC subjects

As our findings uncovered a direct connection between peripheral- and tumor-infiltrating Foxp3E2^+^ Tregs, we assessed whether their peripheral frequency correlated with the clinical parameters of our BC cohorts, which have been stratified based on histopathological analyses (tumor-node-metastasis – TNM) into luminal A and B tumors with different survival periods (*67*). First, we evaluated one of the main prognostic markers in BC, the intratumoral Ki67 expression (*68*). We found that the percentage of intratumoral Ki67 was significantly higher in the high compared to the low E2 PB-ratio BC group (20% *vs* 10%, *P* = 0.022) (**Fig. 8a**). Furthermore, 74% of the BC subjects with low E2 PB-ratio belonged to the luminal A subgroup (which has a better prognosis than luminal B (*67*)), while only 43% of the high E2 PB-ratio BC subjects fell in that subgroup (**Fig. 8b**). Moreover, luminal B BC subjects showed higher PB-ratio compared to the luminal A group and this correlated with stronger suppressive activity (**Fig. 8c, d**). Finally, we stratified our BC cohort in two clinical-pathological groups with different prognosis (*69*) and we found that E2 PB-ratio strictly reflected the overall BC status, as it was significantly increased in the poor-prognosis BC group (0.56 *vs* 0.51) (**Fig. 8e**). Importantly, we show stronger immunosuppression of peripheral Tconvs from Tregs of the poor-prognosis BC group (**Fig. 8f**), thus suggesting that an increased percentage of Foxp3E2^+^ Tregs is associated with an enhanced peripheral suppressive function and worse prognosis. Taken together, our findings identify Foxp3E2 as a novel prognostic marker in BC (**Fig. 8g**).

**Figure 8.**
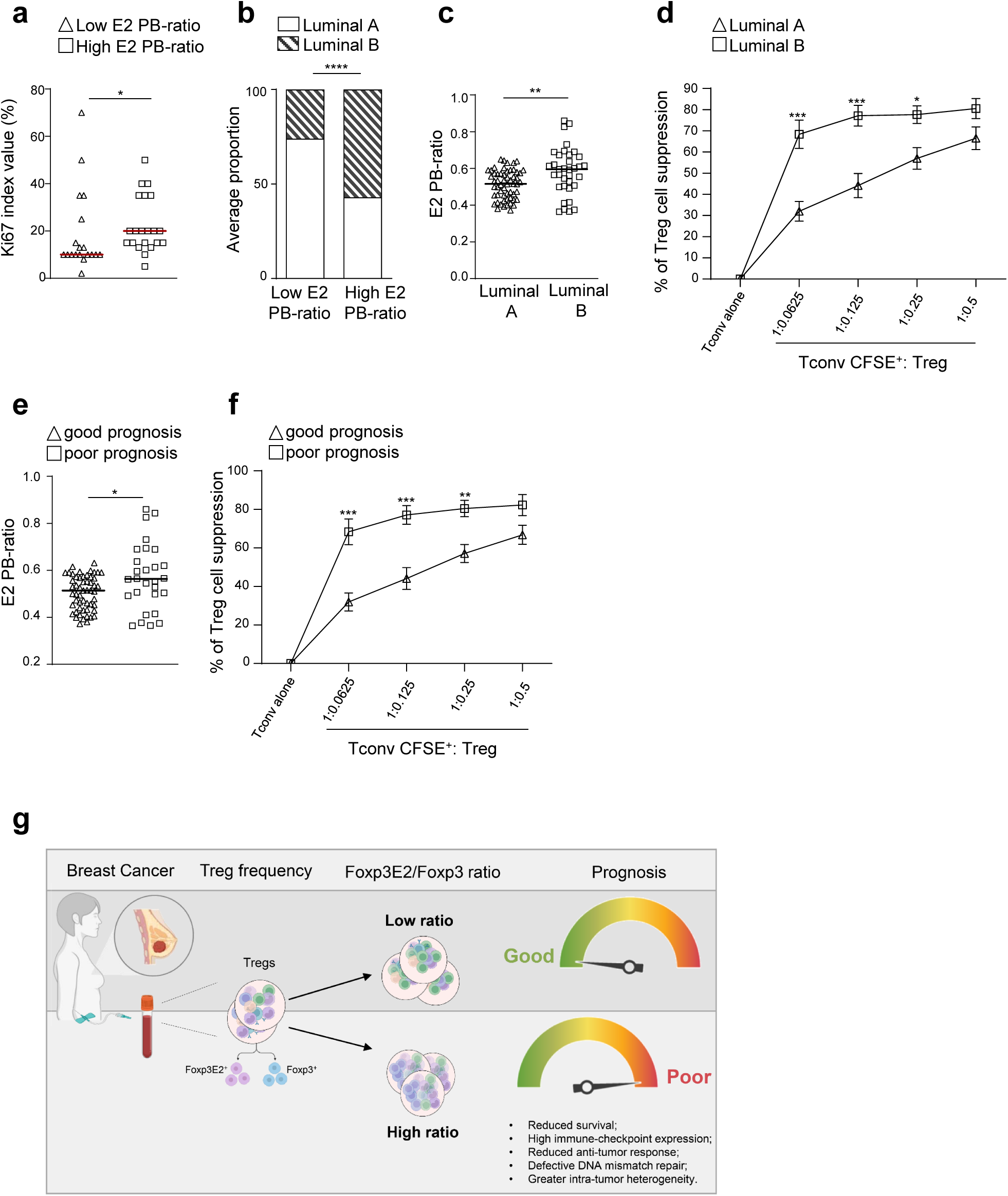
High Foxp3E2^+^/Foxp3^+^ ratio predicts worse prognosis in two independent cohorts of newly diagnosed HR^+^BC subjects. **(a)** Intratumoral Ki67 from BC subjects with high (*n* = 21) and low (*n* =19) E2 PB-ratio. **(b)** Luminal A and B (average proportion) in the high and low E2 PB- ratio BC groups. **(c)** E2 PB-ratio in Luminal B (*n* = 35) and Luminal A (*n* = 57) BC groups. **(d)** Percentage of Treg suppression from Luminal B (*n* = 6) and Luminal A (*n* = 12) BC subjects. **(e)** PB-ratio from BC subjects with poor (*n* = 29) or good (*n* = 54) prognosis. **(f)** Percentage of Treg suppression in BC subjects with poor (*n* = 5) or good (*n* = 13) prognosis. **(g)** Schematical summary of the results. Each symbol shows independent biological samples **(a, b)** or experimental replicates **(c-f)**. Data are presented as Median values. Statistical analyses were performed by using Wilcoxon and Mann-Whitney *U*-test (two tails) **(a, c-f)** and Fisher’s exact test **(b)**; \**P*≤ 0.05; \*\**P*≤ 0.01; \*\*\**P*≤ 0.005; \*\*\*\**P*≤ 0.0001.

## Discussion

Women with early-stage breast cancer (BC) have an independent risk of recurrence and mortality for at least 20 years after the initial diagnosis with the greatest impact demonstrated in hormone receptor-positive (HR^+^) disease, even after 5 years of adjuvant endocrine treatment. In order to improve BC patient survival, an accurate classification of breast cancer subtypes and the identification of prognostic markers that can predict the course of the disease (e.g., relapse, mortality, therapeutic response) are needed alongside with the identification of the underlying molecular pathways.

Regulatory T cells (Tregs) that express the transcription factor Foxp3 are crucial for maintaining immunological *self*-tolerance and suppressing the anti-cancer immune response. The role of distinct Treg subpopulations, their respective functions and interactions within the complex network of the TME have, however, not been fully elucidated (*1*). The composition of intratumoral Foxp3^+^ Tregs is characterized by a subpopulation of highly immune suppressive cells having a distinct gene expression profile, possibly due to the hyperstimulation by tumor-associated antigens (*19, 31, 32*). It has been previously reported that, among all Foxp3^+^ Tregs, those expressing the isoforms retaining exon 2 display stronger suppressive function and increased lineage stability (*34, 37*). Notably, although the role of Foxp3 as a master regulator of Treg differentiation and stability is conserved in mouse (*33, 70*), mouse *Foxp3* gene does not have splicing variants making human studies essential to characterize Tregs and their function in human cancers. To date, whether Foxp3E2^+^ Tregs are preferentially enriched within the TME or the peripheral blood of subjects with cancer and how they correlate with the clinical outcome is completely unknown.

Here, we investigated the role of Tregs in two independent cohorts of newly diagnosed ER^+^PR^+^HER2^-^ (HR)^+^ BC and non-malignant breast fibroadenoma (BF) subjects (i.e., 57 patients). Our analyses revealed for the first time a different composition of tumor-infiltrating immune cells in breast cancer and non-malignant tumors, with BC being characterized by a lower CD8^+^/CD4^+^ ratio and higher frequency of Foxp3^+^ Tregs. Strikingly, we showed that the Foxp3E2^+^ Treg subpopulation is increased in the TME of BC compared to BF patients, and a similar enrichment is detected in the peripheral blood as well. We further associated Foxp3E2^+^ Treg enrichment with worst BC prognosis both in our HR^+^ BC subjects and in a wider published cohort of 990 BC subjects (from the TCGA) that also includes HER2^+^ and triple negative BC.

In addition, we detected a lower frequency of co-mutations in BC subjects with a high Foxp3E2^+^/Foxp3^+^ ratio, suggesting a sub-clonal heterogeneity of these tumors that has already been associated with therapy resistance and tumor shaping (*53–56*). Overall, our data suggest that Foxp3E2 might be used as a novel biomarker to develop a blood-based test predictive of BC prognosis (all tumor subtypes) and, perhaps, of susceptibility to specific therapies.

The origin of intra-tumoral Tregs and their relationship with those circulating in peripheral blood remains unclear. Nonetheless, comparing intra-tumoral Foxp3E2^+^ Tregs with those circulating in the peripheral blood, we found that both overexpress Helios, a transcription factor that regulates Treg function and stability, suggesting that it might be involved specifically in the differentiation/function of the Treg subset expressing the Foxp3E2 splicing variants. Notably, BC tumors with high Foxp3E2^+^/Foxp3^+^ ratio showed greater levels of cancer stem-cell genes (e.g., WNT3a, NANOGP1 and ESRG) suggesting that this Treg subset might shape the TME to foster cancer stem cell growth or maintenance. Importantly, gene expression and mutational signature analyses showed that Foxp3E2 may be utilized to identify a unique subset of individuals with stronger tumor immune tolerance, persistent DNA damage (only ATM-dependent) and metabolic rewiring induced by hyperactive oncogenic signaling (as PI3K/AKT) (*49, 71*). Moreover, intratumor heterogenicity and tumor shaping in the high Foxp3E2^+^/Foxp3^+^ ratio BC group may result from an earlier immunosurveillance that spreads the number of sub-clonal neoantigens associated with increased aggressiveness and drug resistance in cancer (*53, 64, 72, 73*). It is important to note that chemotherapy and radiation-induced mutagenesis may be accelerated in BC patients having a deficiency in the DNA mismatch repair (MMR) system (*74*). Some novel mutated genes may be cancer-driver genes, which means that MMR inactivation can lead to disease progression and therapeutic resistance (*23*). TGCA data mining collectively demonstrate that FOX3E2 expression in the TME is associated with defects in mismatch repair and PI3K/AKT co-mutations indicating stemness and sub-clonal heterogeneity. In addition, FOXP3E2 is associated with immunosuppressive signatures that may contribute to stem-like clones evading tumor immune responses and, account for BC immunological quiescence (low lymphocyte infiltration, low mutational burden, minimal response to immunotherapy (*5, 50, 75*) and tumor shaping (*55*)).

Finally, our data suggest that Foxp3E2^+^ Tregs may account for higher immunosuppressive function. Strikingly, we showed that Tregs from BC patients with increased Foxp3E2 levels provide stronger suppression of effector cells during *in vitro* functional assays. This is consistent with the increased expression of Helios, ICOS and immune checkpoint receptors by Treg immunophenotyping and with the enrichment of an immunosuppressive gene signature in the BC patients with high Foxp3E2^+^/Foxp3^+^ ratio. In addition, known markers of anti-tumor T cell response (e.g., CD8^+^/Treg ratio or CCR8 expression) strongly suggest that this Treg subset is associated with tumor immune escaping. Of note, the increased expression of ICOS and lower levels of pS6 suggest that Treg function might be tuned by the mTOR metabolic pathway.

Our overall data suggest that the Foxp3E2^+^ Treg subpopulation might have a dominant role in cancer evasion/immunosuppression. This might at least in part be mediated by the increased expression of immune checkpoint co-stimulatory receptors (e.g. CTLA4, PD1 and TIGIT), which represent the targets of currently available immunotherapies that are effective against several malignancies, including BC (e.g., ipilimumab and pembrolizumab) (*1, 76–80*). Notably, the levels of CCR8 are also increased in TIL-Foxp3E2^+^ Tregs. As elevated CCR8 levels in TIL-Tregs associate with poor prognosis in several cancer types (*19, 66*), this could further contribute to Foxp3E2^+^ Treg retention within the tumor thus amplifying the immunosuppression. Altogether, these highlight the importance to consider TIL-Foxp3E2^+^ Tregs as a novel target for improving the actual BC immunotherapeutic strategies. We hypothesized that breast cancer cells may induce (and/or increase recruitment/retention) of Foxp3E2^+^ Tregs through the establishment of a highly immunosuppressive milieu (**Fig. S10b**). To this aim, unravelling the specific pathways involved in Foxp3E2^+^ Treg induction will be instrumental to restrain their generation restoring the tumor- immune responses.

In conclusion, we showed that high Foxp3E2^+^/Foxp3^+^ (E2) ratio in the peripheral blood of BC subjects reflects stronger immunosuppression and defective MMR at the tumor site, thus predicting poor prognosis. Since the accumulation of Tregs represents an essential mechanism for cancer immune evasion and a critical barrier to anti-tumor immunity and immunotherapy, our findings may represent a vital jigsaw piece in the early detection of the BC prognosis puzzle. Furthermore, these results might offer a novel paradigm for developing a “super-targeted” approach that selectively restrains tumor-promoting Tregs while preserving a proper peripheral tolerance.

## Methods

### Subjects and study design

The clinical and demographic characteristics of the study cohorts were shown in **Supplementary Table 1**. Female subjects were enrolled after obtaining informed consent. The study has been approved by the Institutional Review Board of the University of Naples “Federico II” (Protocol n. 269/15/ES01). Biological samples were collected by clinicians at the National Cancer Institute – IRCCS “G. Pascale” Foundation (Clinical and Experimental or Oncological Surgery of Senology) and at the Department of General, Oncological, Bariatric and Endocrine-Metabolic Surgery, University of Naples “Federico II”, Naples. BC subjects were naïve- to-treatment and with definite clinicopathological parameters, including age, tumor-node- metastasis (TNM) stage, histological type and grade (according to WHO 2012-2019 and Elston- Ellis) (*81, 82*), Ki67 index, estrogen receptor (ER), progesterone receptor (PR) and human epidermal growth factor receptor 2 (HER2) status. For each subject, a detailed past medical history was recorded to exclude intake of glucocorticoids and/or antihistamine drugs in the 2 months preceding the enrolment and previous diagnosis of chronic inflammatory, autoimmune or other neoplastic diseases. Subjects underwent breast surgery or core needle biopsies, collected with ultrasound guidance. Tissue and blood samples were collected prior to chemotherapy, radiotherapy, endocrine therapy, or any other treatment. Enrolled subjects were classified into immunohistochemically defined surrogate molecular subtypes, according to the American Society of Clinical Oncology/College of American Pathologists (ASCO/CAP) 2013-2018. Healthy female donors (HD) were matched for age and body mass index and had no history of inflammation, endocrine or autoimmune disease. The ethnic distribution among the groups was comparable, with all participants being white.

### Breast cancer and breast fibroadenoma tissue samples preparation

For the preparation of tissue microarray (TMA) and histologic review, five-micrometer sections from each formalin-fixed-paraffin-embedded tissue block were stained with hematoxylin and eosin for the identification of tumor areas. Immunohistochemical (IHC) analysis was performed on TMA with up to four 1.5-mm cores from primarily the invasive tumor front from each tumor. A review of histologic subtype and grade was performed according to WHO guidelines (*81*). The diagnosis of ductal carcinoma with medullary characteristics was designated for high-grade tumours with pushing margins and syncytial growth patterns in >75% of the tumour in association with a pronounced lymphoplasmacytic infiltrate (*81*).

### Immunohistochemistry

Immunohistochemical staining was performed on slides from formalin-fixed, paraffin-embedded tissues to evaluate the expression of CD3, CD8, Foxp3 and Foxp3E2 markers in breast fibroadenoma (*n* = 6) and breast cancer (*n* = 23) tissues. Paraffin slides were de- paraffinized in xylene and rehydrated through graded alcohols. Antigen retrieval was performed with slides heated in 0.01 M citrate buffer (pH 6.0) in a bath for 20 minutes at 97°C. After antigen retrieval, the slides allow to cool. The endogenous peroxidase was inactivated with 3% hydrogen peroxide was inactivated with 3% hydrogen peroxide. After protein block (BSA 5% in PBS 1x), slides were incubated with specific primary antibodies: human anti-CD3 (2GV6) dilution 1:100 (Ventana), human anti-CD8 (CAL66) dilution 1:100 (Roche), human anti-FOXP3 (D2W8E) dilution 1:125 (Cell Signaling) and anti-human Foxp3E2 (150D) dilution 1:125 (BioLegend). The sections were incubated for 1 hour with Novocastra Biotinylated Secondary Antibody (HRP-conjugated) and visualized with 3,3’-Diaminobenzidine (DAB) chromogen. Finally, the sections were counterstained with hematoxylin and mounted. CD3, CD8, Foxp3 and Foxp3E2 positive nuclei were counted evaluating at least five fields at 400x magnification. All sections were evaluated in a blinded fashion by 2 investigators. For each marker, a mean value of up to five cores for each patient was calculated representing the overall expression of the specific marker.

### Breast tissue preparation and cell purification

For the isolation of tumor-infiltrating lymphocytes (TILs), dissected tissue fragments from freshly isolated biopsies were transferred in GentleMACS C tubes (Miltenyi Biotec) containing Hanks’ Balanced Salts Solution (HBSS) with Calcium, Magnesium, Sodium Bicarbonate and without Phenol Red (Aurogene) supplemented with 0.5 mg/mL Collagenase IV (Sigma), 50 ng/mL DNAse I (Worthington), 2% fetal bovine serum (FBS) (GIBCO) and 10% bovine serum albumin (Sigma). Tissue dissociation was made on a GentleMACS Dissociator (Miltenyi Biotec) by using the “h_tumor 01_03” program. Single-cell suspensions were obtained by disrupting the fragments with a syringe plunger over a cell strainer (100 μm) and washing with cold HBSS. Cell suspension was centrifuged at 2700 rpm for 5 minutes to remove debris and the cell pellet was resuspended in RPMI 1640 medium for successive evaluations. Peripheral blood mononuclear cells (PBMCs) from BF, BC and HD subjects were isolated from blood samples after Ficoll-Hypaque gradient centrifugation (GE Healthcare). Tregs (CD4^+^CD25^+^CD127^-^) and Tconvs (CD4^+^CD25^-^) were purified (90-95% pure) by using the CD4^+^CD25^+^ Regulatory T Cell Isolation Kit (Miltenyi Biotec).

### Flow cytometry, proliferation and CFSE staining

Freshly isolated PBMCs and TILs from BF, BC and HD females were surface-stained with the following mAbs: APC-H7-conjugated anti-human CD45 (2D1), V500–conjugated anti-human CD4 (RPA-T4), APC-H7–conjugated anti-human CD4 (RPA-T4), PE-Cy7–conjugated anti-human CD8 (RPA-T8), BV421–conjugated anti-human CD279/PD-1 (EH12.1) and BV421–conjugated anti-human CD198/CCR8 (4333H) all from BD Biosciences, PE-Cy7–conjugated anti-human TIGIT (MBCA43) (eBioscience). Thereafter, cells were washed, fixed and permeabilized (anti-human FOXP3 staining Set PE; eBioscience) and stained with following mAbs: PE-conjugated anti-human FOXP3 from eBioscience (PCH101, that recognizes all splicing variants through an epitope of the amino terminus of Foxp3), and PE-conjugated anti-human Foxp3 from eBioscience (150D/E4, that recognizes Foxp3E2 variants through an epitope present in the exon 2 only), APC–conjugated anti-human CD152/CTLA-4 (BNI3) (BD Biosciences), Alexa Fluor 488–conjugated anti-human Helios (22F6) and BV510– conjugated anti-human Ki67 (B56). Cells were analyzed with FACSCanto II (BD Biosciences) and FlowJo software (Tree Star). For T cell proliferation and suppression assays, Tconv cells (2 × 10^4^ cells/well) were stained with the fluorescent dye CFSE at 1 μg/ml (Invitrogen). Flow cytometry analyzing CFSE dilution was performed by gating on CFSE^+^ cells stimulated for 72 hours in round-bottomed 96-well plates (Corning Falcon) with anti-CD3/anti-CD28 mAb-coated beads (0.2 beads/cell; Thermo-Fisher) alone or cultured with Tregs from BC and HD subjects, respectively.

### Systematic transcript variant analysis in public databases

The Foxp3 spliced variant sequences were assessed in UCSC Genome Browser on Human (GRCh37/hg19) (*83*) databases. The schematic diagram of the Foxp3 variant structures is reported in **Fig. S1d**.

### The cancer genome atlas (TCGA) BRCA database analyses

Foxp3 splicing variant expression data derived from TCGA splicing variant database (TSVdb) web tool (http://www.tsvdb.com) and are reported as normalized RNA-Seq by Expectation Maximization (RSEM) values. Samples with unreported and/or missing clinical data were removed. The Foxp3E2^+^/Foxp3^+^ ratio was calculated using GRCh37/hg19 coordinates chrX:49,114,121-49,114,225 and chrX:49,109,587-49,109,663 that recognize respectively the Foxp3 splicing variants containing the exon 2 and the exon 9 (being this last common to all transcripts).

### Kaplan–Meier survival plot

Overall survival analysis was conducted using only patients with survival and gene expression data from TSVdb. Samples were categorized using Cox proportional hazards regression into two groups based either on the mean RSEM value (high expression ≥ Q2 and low expression < Q2) or on the upper quartile RSEM value (high expression ≥ Q3 and low expression < Q3). The Kaplan–Meier survival plots were generated using R packages: “survival and survminer”. The survival curves of samples with high and low gene expression were compared by log-rank test, and data groups with *P* value < 0.05 were considered statistically significant.

### RNAseq analysis

Primary BC (*n* = 990) and normal breast tissue (*n* = 112) RNA-seq data counts were downloaded from the TCGA BRCA project (available online at https://portal.gdc.cancer.gov/projects/TCGA-BRCA). Further analysis and visualizations of the processed data were performed in R and Bioconductor. For differential expression analyses between high- (*n* = 248) and low- (*n* = 742) ratio BC groups, counts were normalized using the size factor normalization technique available in DESeq2 and an absolute log2FoldChange > 0.5 and p- adj < 0.001. We used the online tools RDAVIDWebService (*84*) and GOplot (*85*) to identify GO Biological Processes overrepresented and to prepare circular composition overview. We performed a statistical overrepresentation test using default parameters. GO-terms were considered overrepresented only if FDR-corrected P-values were below < 0.05. Then, ClusterProfiler v.4.6 and Enrichplot v.1.19.0.01 were used for gene set enrichment analysis (GSEA) and plotting (*49, 86*). DNA mismatch repair (MMR) gene expressions were obtained by comparing the low- (*n* = 742) and high- (*n* = 248) ratio BC groups and filtering the normalized count matrix.

### Tumor immune microenvironment cell composition analysis

Tissue composition analysis of low (n = 735) and high (n = 248) ratio BC immune and stroma (Tumor immune microenvironment deconvolution) was performed using the online tool TimeDB (*63*) based on differentially expressed genes (DEGs) obtained from RNAseq analysis.

### Mutation enrichment analysis

Variants were obtained from TCGA-BRCA WES using the TCGAbiolinks R package (*87*) to identify differentially mutated genes in low- (*n* = 650) and high- (*n* = 211) ratio BC groups. Analysis (variants number, somatic interactions, APOBEC enrichments, and signatures detection) and visualization of mutations were performed using the Maftools R package (*88*). Contributions of mutational signatures in COSMIC(*89*) were determined in each sample using nonnegative matrix factorization provided by the NMF v1.8.0 R package (*90*) using a p-value < 0.001

### Differentially methylated regions

Differentially methylated regions were calculated using the normalized beta-values (methylation values ranging from 0.0 to 1.0) obtained from TCGA-BRCA Illumina Human Methylation 450 downloaded through TCGAbiolinks R package. To compare low- (*n* = 524) and high- (*n* = 194) ratio BC groups, we used the Wilcoxon test with the adjusted Benjamini-Hochberg method. The default parameters were set to require a minimum absolute beta- value difference of 0.2 and a p-value adjusted of < 0.01.

### RPPA analysis

Proteomic analyses were performed using the level 4 (log2 transformed with loading and batch corrected) RPPA dataset from the TCGA-BRCA study downloaded from The Cancer Proteome Atlas portal (https://tcpaportal.org/tcpa/). For differential protein expression analysis between high- (*n* = 248) and low- (*n* = 742) ratio BC groups, RPPA relative fluorescence intensity (RFI) values were compared using an ANOVA FDR p-value threshold of less than 0.05. The data were then scaled based on Average RFI threshold for each protein to extract upregulated (red) and downregulated (green) probes of the high-ratio BC group. Metascape was used to perform the enrichment analysis of the differentially expressed probes. The data were displayed as median values.

## Statistical analysis

Statistical analyses were performed using GraphPad program (Abacus Concepts) and R packages. Results were expressed as Median and interquartile range (IQR). The non-parametric Mann-Whitney *U*-test, the Wilcoxon matched-pairs signed-rank test and the t-test were used. Correlations were computed with a non-parametric Spearman r correlation test, overall survival with a log-rank test, and hazard ratio with multivariate Cox regression model reference. A two-tailed *P* value < 0.05 was considered statistically significant.

## Data and code availability statement

The results published here are based in part on data from The Cancer Genome Atlas pilot project established by NCI and the National Human Genome Research Institute. The data was retrieved partly via public repositories and partly via the Genotypes and Phenotypes Authorization Database (dbGaP) (accession number: phs000178.v11.p8). Links to public repositories can be found via online citations. Analysis and visualizations on the processed data were performed using citated R packages. Clinical datasets that support the findings of this study are not publicly available due to information that could compromise research participant consent. Each request for access to the dataset will be granted upon reasonable request sent to the corresponding author and approval by the ethic committee.

## Acknowledgments

We thank the study participants of this study, Mariarosaria Montagna e Salvatore De Simone for technical support, Dr. Gjada Criscuolo and Dr. Remo Poto for critical reading of the manuscript. IC acknowledges support by FISM - Fondazione Italiana Sclerosi Multipla cod.2020/BC/001 and financed or co-financed with the "5 per mille" public funding. AL acknowledges support by Fondazione Umberto Veronesi.

## Author contributions

CF, FDR and VDR contributed to study design. FDR, VG, FC, EE, ID, LI, AA, MP, FP, FG, DR, MM, BZ and MdB contributed to the collection of clinical specimens. FDR, AA and LI provided clinical data. CF, AC, AL, ALF, TM and AF performed laboratory experiments. CF, AL, ALF and BDS collected the data. APezone and AP performed NGS data analysis. GTM, AS, GV and GM reviewed and edited the draft. VDR, IC and APezone drafted the manuscript.

## Competing Interests Statement

The authors declare no competing interests.

## Funding

This work was supported by grants from Fondazione Italiana Sclerosi Multipla (2018/R/4), Ministry of Education, University and Research (MIUR) PRIN 2022KT2HBJ, PRIN- PNRR 2022C5KBT, European Union - Next Generation EU “PE8 Ageing Well in an ageing society – AGE-IT” Investment 1.3 (Partenariato Esteso - PE0000015) and Associazione Italiana per la Ricerca sul Cancro-TRIDEO (Transforming Ideas in Oncological research, n.17447) to VDR.

---

**Supplementary Figure 1.**
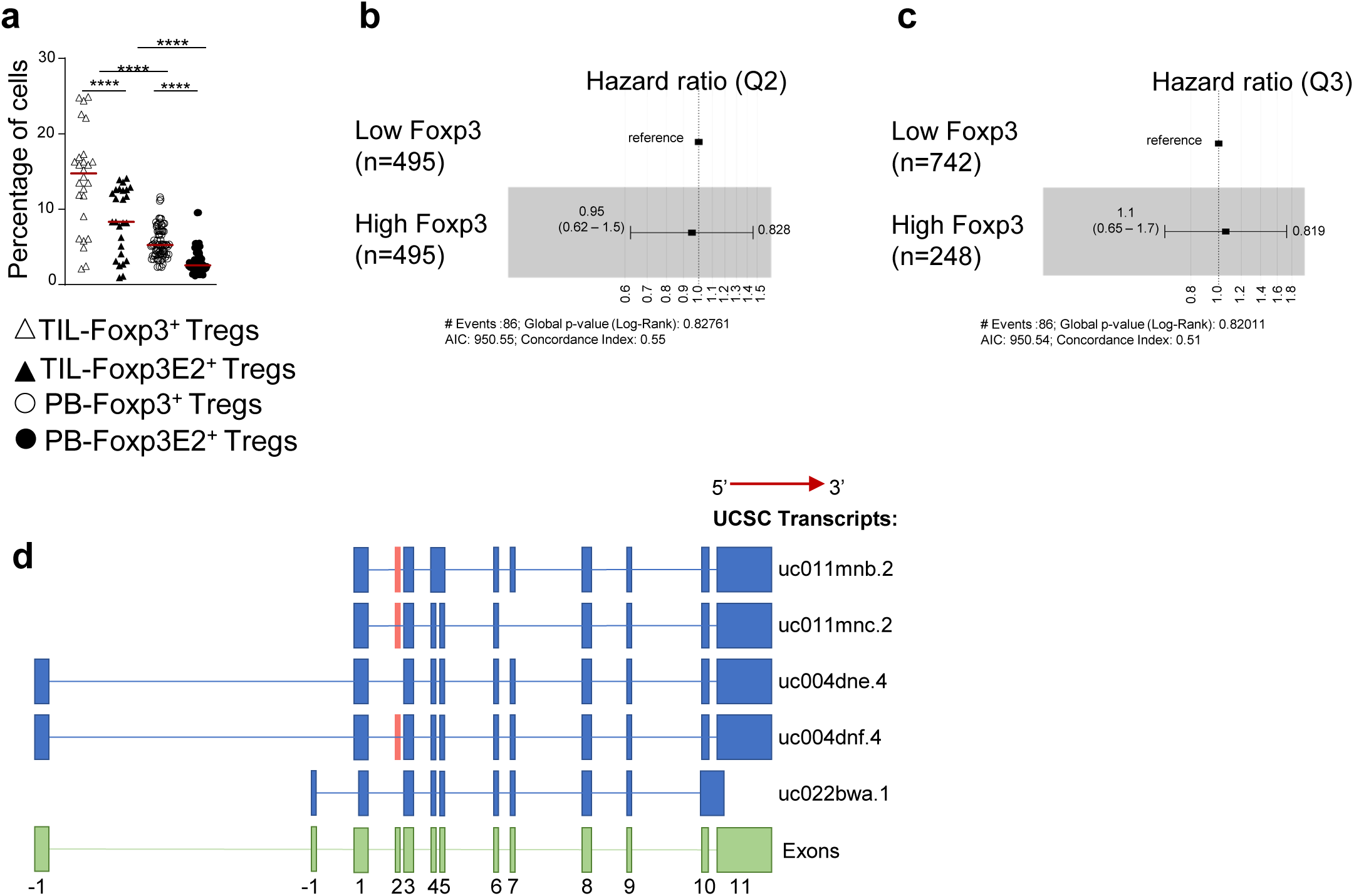
RNAseq analysis of Foxp3 transcript expression in primary tumor of breast cancer (BC) subjects. **(a)** Percentage of Foxp3^+^ and Foxp3E2^+^cells (gated on CD4^+^) from TILs [respectively empty and full triangles (*n* = 26)] and PB [respectively empty and full dots (*n* = 77)] of BC subjects. Each symbol showed experimental replicates. **(b)** Hazard ratio (HR) (0.95, CI 0.62 – 1.5, Cox *P* = 0.828) of BC subjects (*n* = 990) stratified into low- (*n* = 495) and high- (*n* = 495) Foxp3 expression based on its median value, obtained through RNAseq data using TSVdb. **(c)** HR (1.1, CI 0.65 – 1.7, Cox *P* = 0.819). **(d)** Schematic representation of the five alternative Foxp3 transcripts: the exon 9 is common to all mRNAs, while only the uc011mnb.2, uc011mnc.2, uc004dnf.4 variants contain the exon 2 (light red rectangle). Data are presented as Median values. Statistical analyses were performed by using Wilcoxon and Mann-Whitney *U*-test (two tails) **(a)**, log- rank test **(b, c)**. \*\*\*\**P*≤ 0.0001.

**Supplementary Figure 2.**
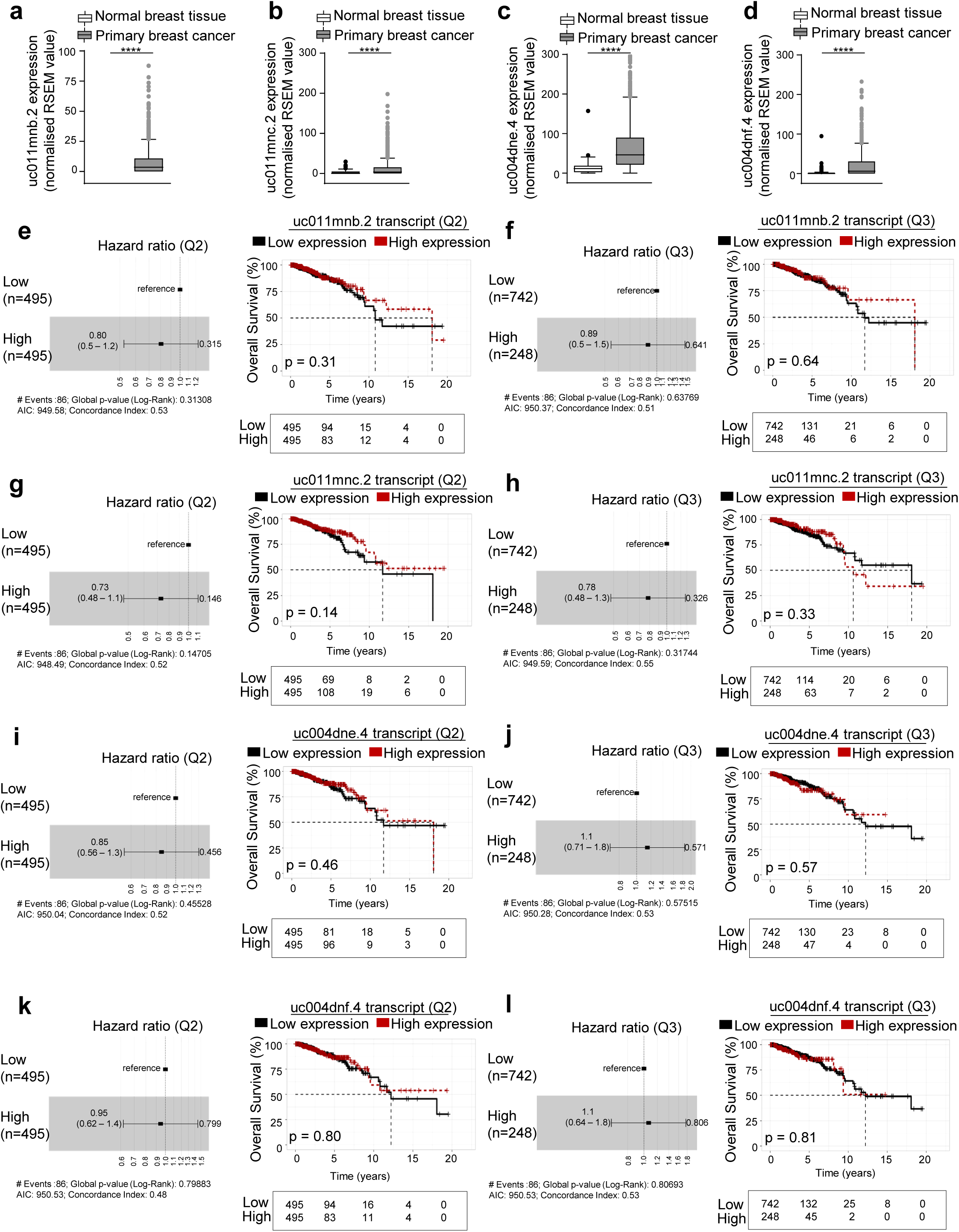
Foxp3 splicing variant expression in normal breast *versus* breast cancer tissue. **(a)** uc011mnb.2, **(b)** uc011mnc.2, **(c)** uc004dne.4, **(d)** uc004dnf.4 Foxp3 transcript expression (normalized RNA-Seq by Expectation Maximization (RSEM) value) in normal breast (*n* = 112) and primary breast cancer (*n* = 990) tissue from TSVdb. **(e-l)** HR and KM survival curves of BC subjects stratified into low- and high-transcript expression based either on the median **(e, g, i, k)** or on the upper quartile value **(f, h, j, l)** of **(a)** uc011mnb.2 **(e, f)**, **(b)** uc011mnc.2 **(g, h)**, **(c)** uc004dne.4 **(i, j)** and **(d)** uc004dnf.4 **(k, l)** expression in primary BC tissue. Data are presented as Median values **(a-d)**. Statistical analyses were performed by using Mann-Whitney *U*-test (two tails) **(a-d),** log-rank test and Multivariate Cox regression model reference **(e-l)**. \*\*\*\**P*≤ 0.0001.

**Supplementary Figure 3.**
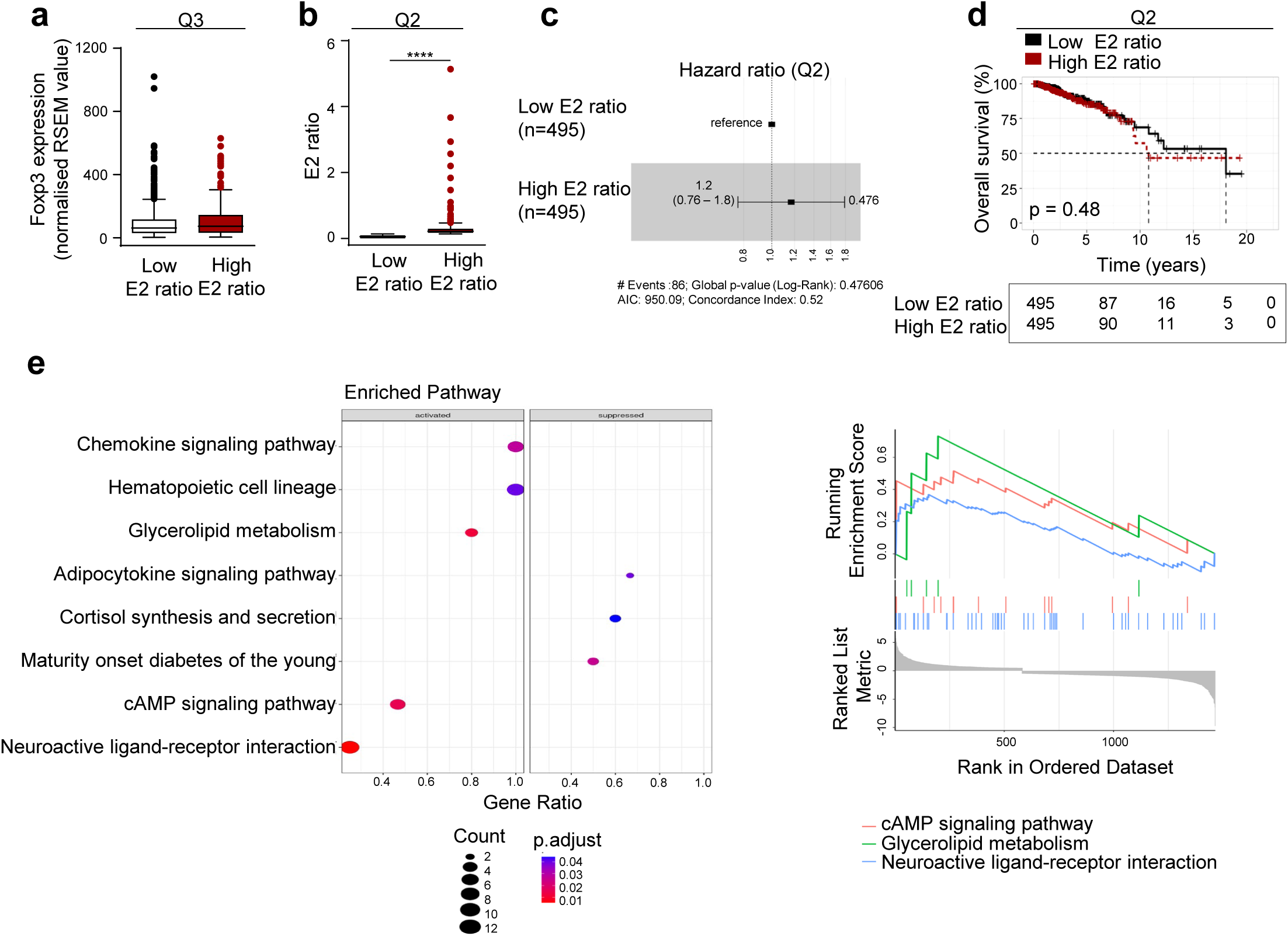
Stratification of BC subjects according to the Foxp3E2^+^/Foxp3^+^ ratio. **(a)** Foxp3 expression levels in low- (n = 741) and high-E2 ratio (n = 249) BC subjects according to the Q3 value cut-off. **(b)** E2 ratio from BC subjects stratified into low (*n* = 495) and high (*n* = 495) according to its median value. **(c)** HR (1.2, CI 0.76 – 1.8, Cox *P* = 0.476) and **(d)** KM survival curve of low- and high- E2 ratio BC subjects. **(e)** Enrichment of KEGG pathways within activated and suppressed genes in high- *vs* low-E2 ratio BC groups. Upper: Significantly enriched KEGG pathways among up- and down- regulated DEGs. The plot shows KEGG terms (vertical axis) versus gene ratio (horizontal). Number of genes in each category is proportional to circle value and color represents the adjusted p-value. Lower: GSEA-based KEGG-enrichment plots of representative gene sets from activated paths: cAMP signaling pathway, glycerolipid metabolism and neuroactive ligand-receptor interaction. Statistical analyses were performed using Mann-Whitney *U*-test (two tails) **(a, b)**, log- rank test **(c)**, Multivariate Cox regression model reference **(d)**. \*\*\*\**P*≤ 0.0001.

**Supplementary Figure 4.**
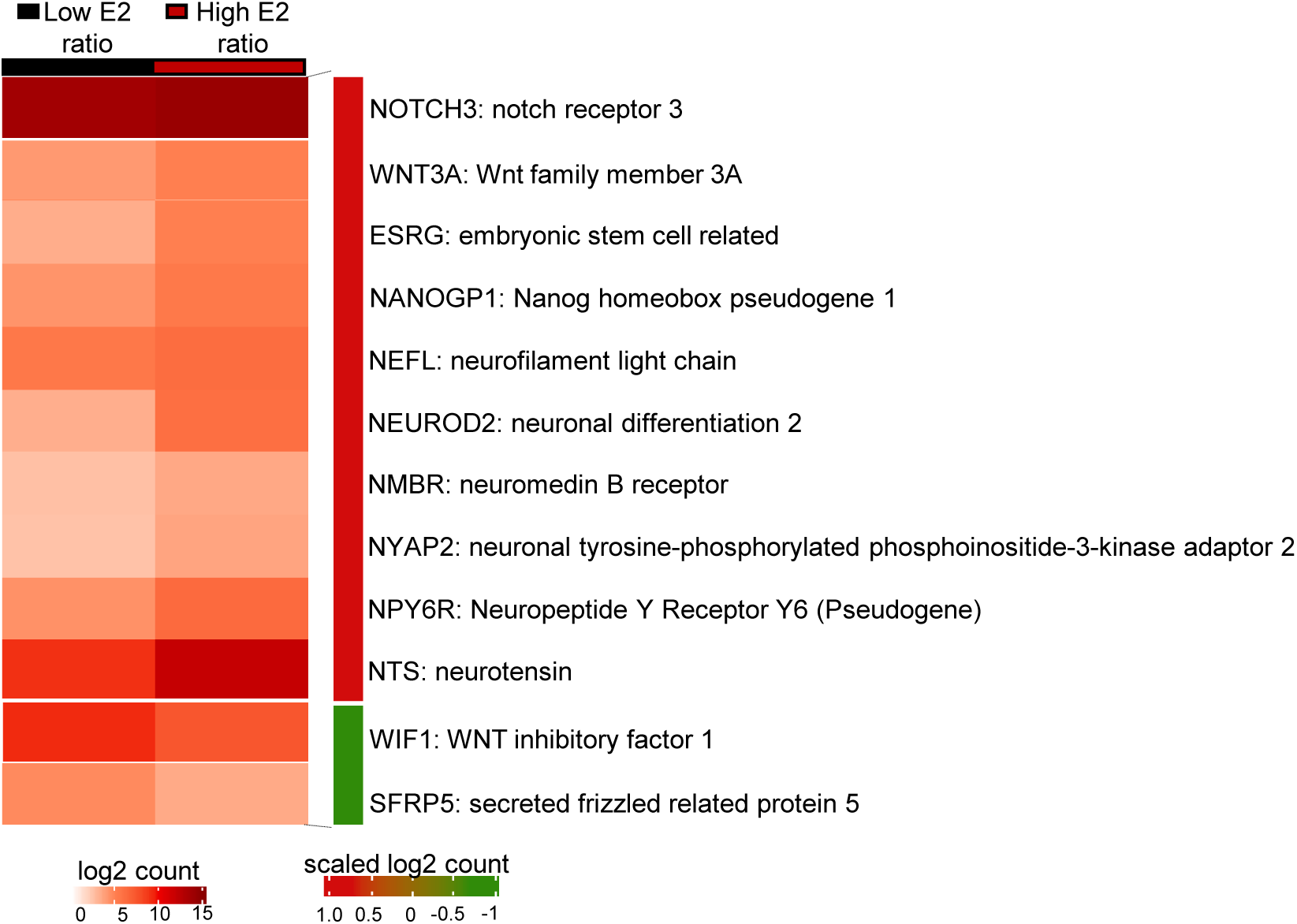
Expression of stem cell-like features in BC subjects according to the Foxp3E2+/Foxp3+ ratio. Heatmap of 12 differentially expressed genes indicating stem cell-like characteristics. Each row corresponds to one gene, while each column represents one group. In the high-E2 ratio BC group, a scaled log2 count revealed that 10 genes were upregulated (red bar) and 2 were downregulated (green bar).

**Supplementary Figure 5.**
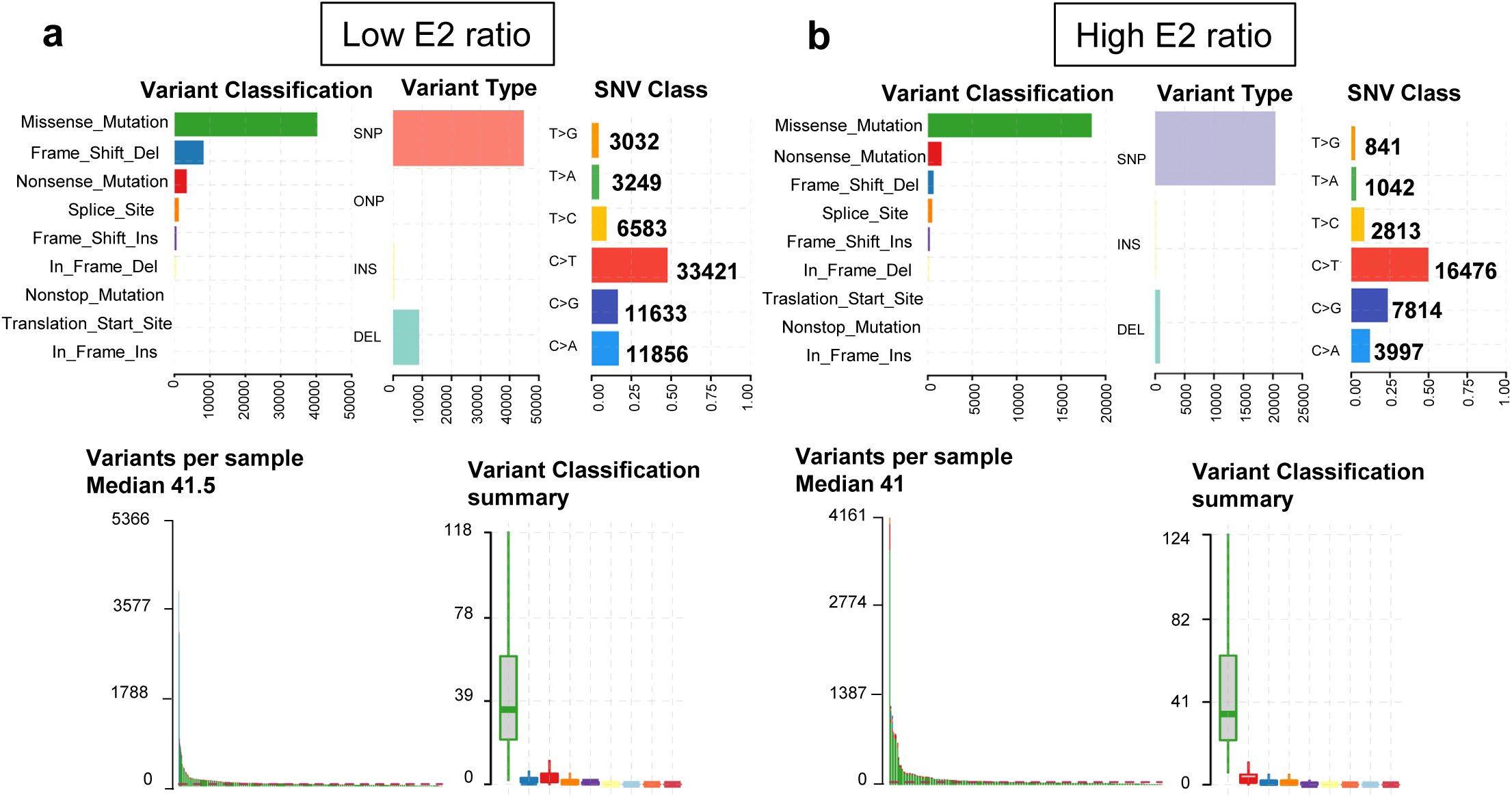
Detection of somatic alterations and association with Foxp3E2+/Foxp3+ ratio. (a,c) Summary of somatic variants displaying variant classification, types, SNV class and number in BC subjects with **(a)** low- and **(b)** high-E2 ratio.

**Supplementary Figure 6.**
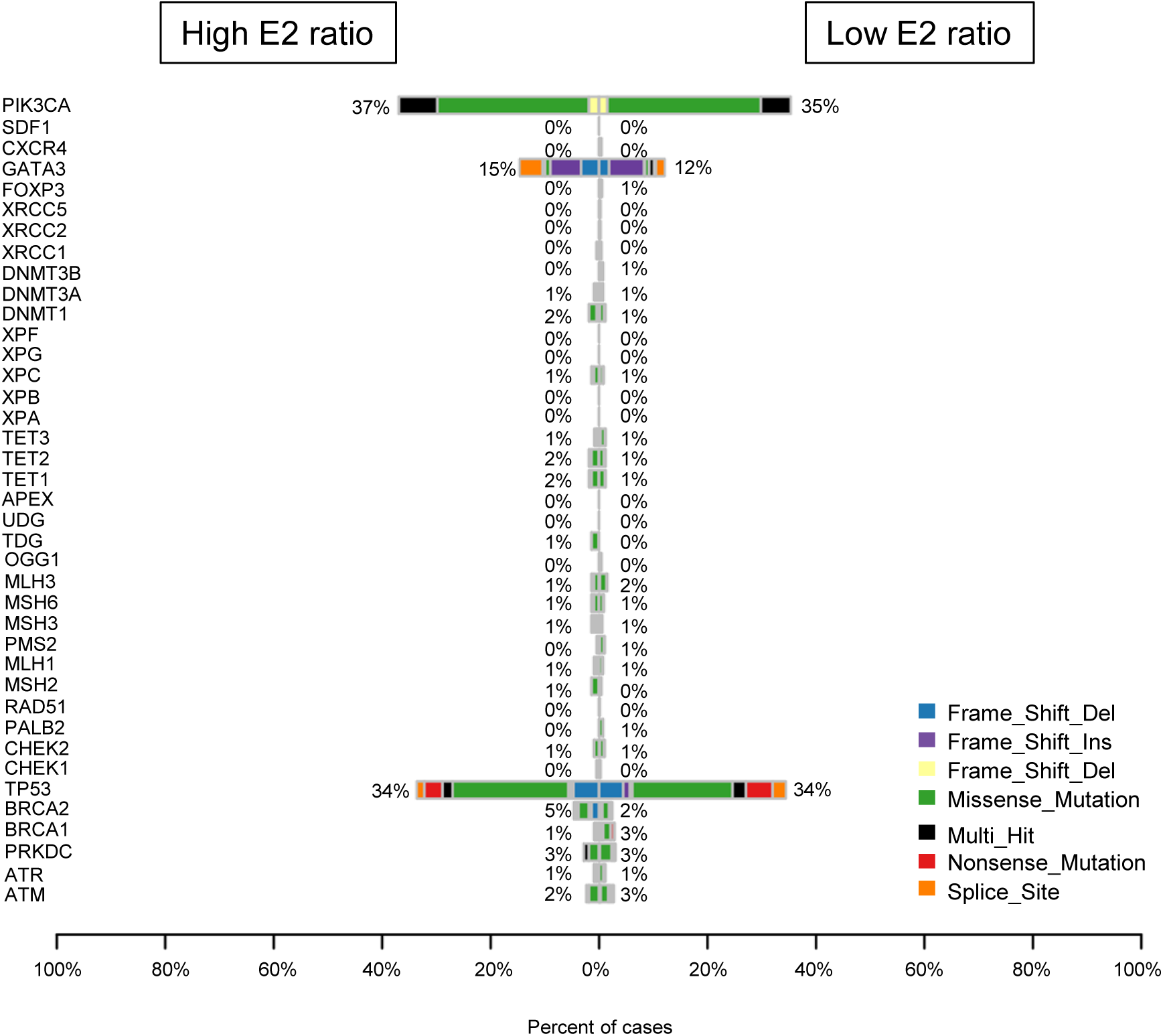
CoBarplot of DNA Damage Repair (DDR) genes. Mutation of DDR genes in high- (left, *n* = 211) and low- (right, *n* = 650) E2 ratio BC groups. No differences were observed in variant incidence between the two groups, although they showed higher occurrences of TP53 mutations. Variant type is indicated (see key below). Data were analyzed using Maftool R package.

**Supplementary Figure 7.**
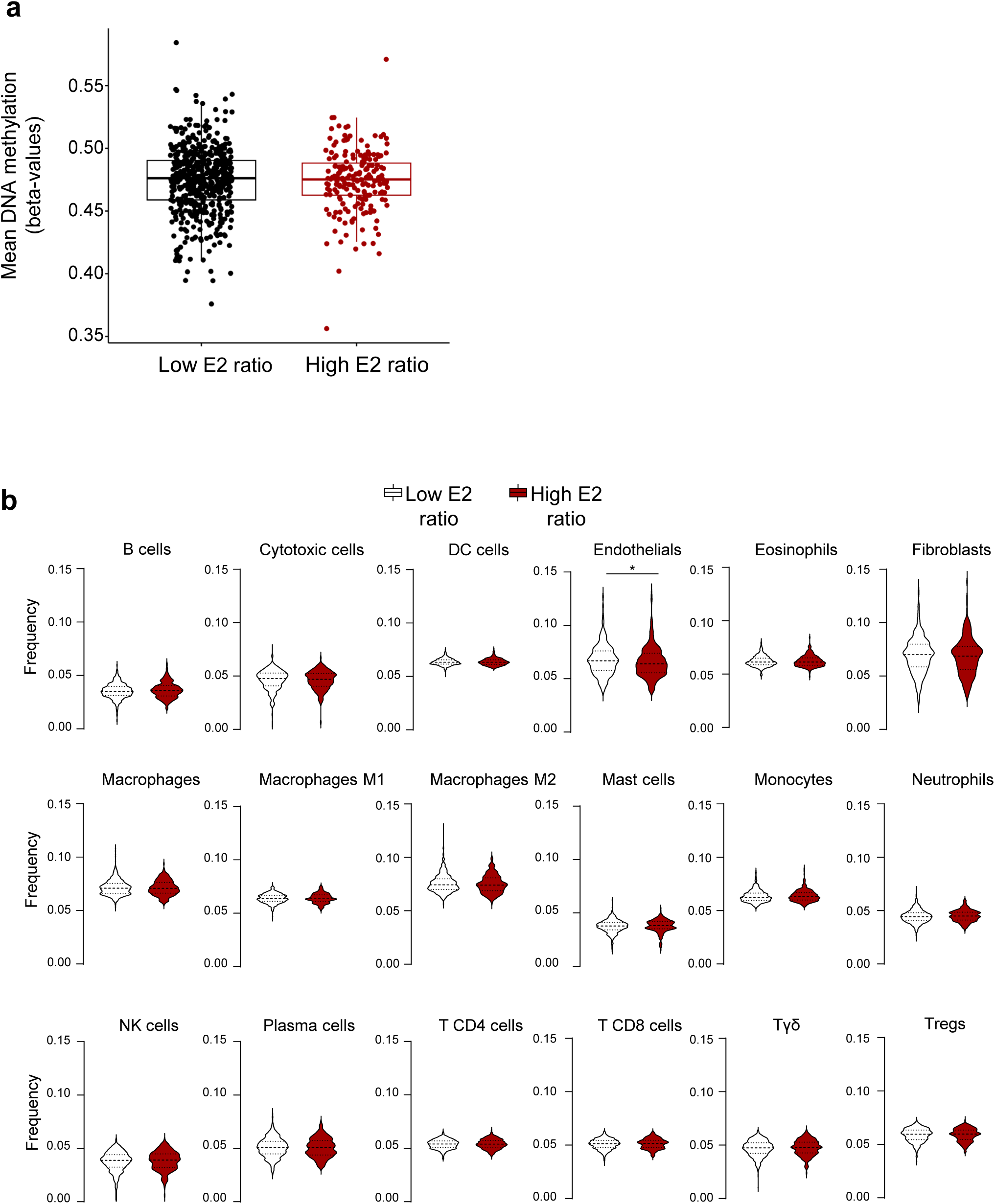
Global DNA methylation analysis and Tumor immune microenvironment deconvolution of primary BC tissues. **(a)** Total mean DNA methylation values for low- (*n* = 524) and high-(*n* = 194) E2 ratio BC groups were calculated by mean methylation beta values (ratio of intensities between methylated and unmethylated alleles) for all probes in the Illumina 450k methylation array TCGA BRCA. **(b)** Violin plots showing the distribution of cell type fraction in primary breast cancer tissue (Tumor immune microenvironment deconvolution) from BC subjects stratified into low- (*n* = 735) and high- (*n* = 248) E2 ratio according to its Q3 value. Tumor immune microenvironment deconvolution was performed using the online tool TimeDB (*62*). Data are presented as Median values. Statistical analyses were performed using Mann-Whitney *U*-test and Wilcoxon signed-rank test (two tails). \**P*≤ 0.05.

**Supplementary Figure 8.**
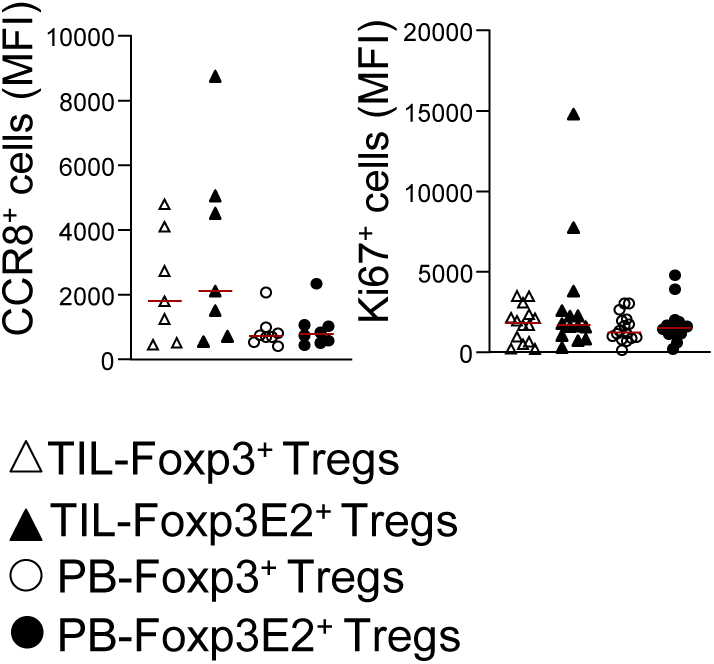
Immunophenotype in Tregs from TIL and PB of newly diagnosed HR^+^ BC subjects. Cumulative data of flow cytometry analysis showing mean fluorescence intensity (MFI) of CCR8^+^ and Ki67^+^ cells (gated on CD4^+^Foxp3^+^ and CD4^+^Foxp3E2^+^) in freshly isolated TILs (at least *n* = 6) and PB (at least *n* = 10) from BC subjects. Each symbol shows independent biological samples. Data are presented as Median values. Statistical analysis was performed by using Wilcoxon and Mann-Whitney *U*-test (two tails).

**Supplementary Figure 9.**
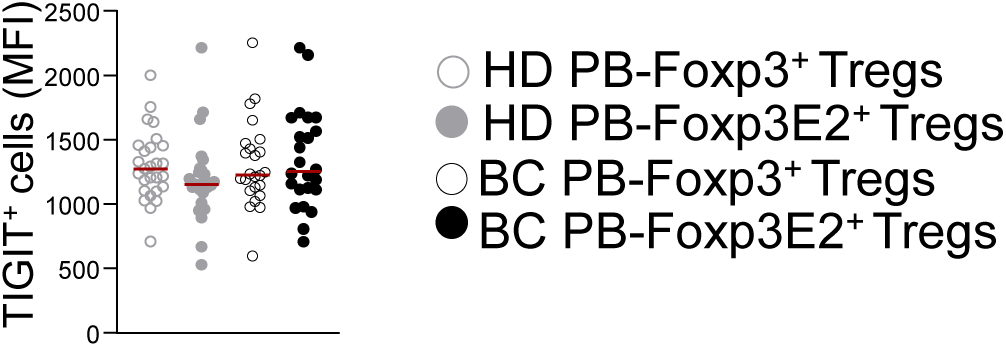
Immune checkpoint expression in Tregs from BC and Healthy Donor (HD) subjects. Cumulative data of flow cytometry analysis showing mean fluorescence intensity (MFI) of TIGIT^+^ cells gated on CD4^+^Foxp3^+^ and CD4^+^Foxp3E2^+^ Tregs from freshly isolated PB of HD (*n* =27) and BC (*n* = 24) subjects. Each symbol shows independent biological samples. Data are presented as Median values. Statistical analysis was performed by using Wilcoxon and Mann-Whitney *U*-test (two tails).

**Supplementary Figure 10.**
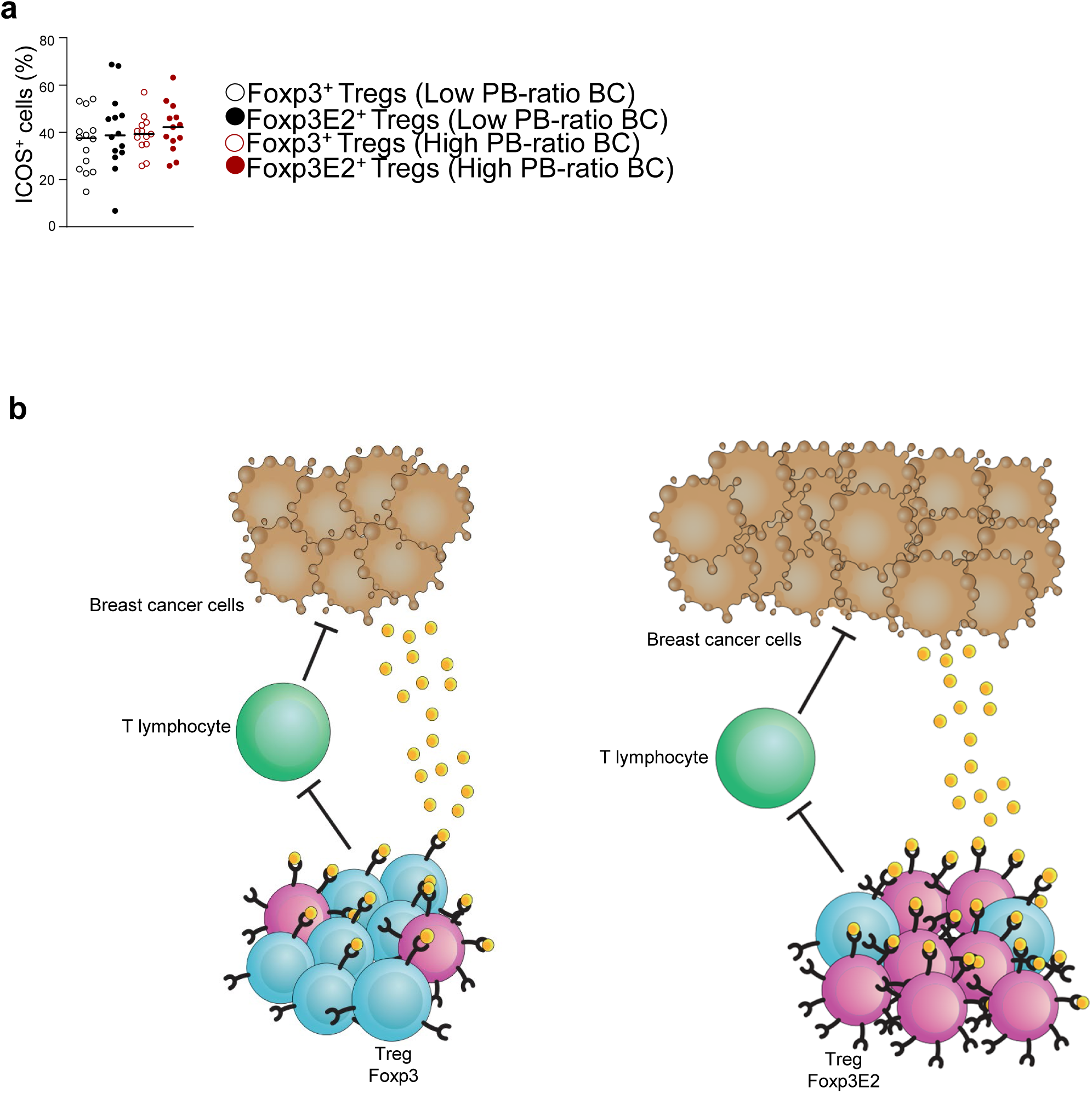
Immunophenotype in Tregs from PB of high- and low-E2 PB-ratio HR^+^ BC subjects. **(a)** Cumulative data calculated by flow cytometry quantification showing the percentage of ICOS^+^ cells gated on CD4^+^Foxp3^+^ and CD4^+^Foxp3E2^+^ Tregs from peripheral blood of high- (*n* = 13) and low- (*n* = 14) PB E2 ratio BC subjects. **(b)** Schematical model of the breast cancer- induced polarization of Foxp3E2^+^ Tregs in the tumor microenvironment. Each symbol shows independent biological samples. Data are presented as Median values. Statistical analysis was performed by using Wilcoxon and Mann-Whitney *U*-test (two tails).

**Supplementary Table 1.** Clinical and demographic characteristics of the study cohort.

|  | **BC** | **BF** | **HD** |
| --- | --- | --- | --- |
| **N <sup>(a)</sup>** | 40 | 17 | 38 |
| **Gender n (%)** |  |  |  |
| Female | 40 (100) | 17 (100) | 38 (100) |
| Male | 0 (0) | 0 (0) | 0 (0) |
| **Age, mean ± SD, year** | 53 (±12) | 39 (±10) | 47 (±10) |
| **Tumor staging n (%)** |  |  |  |
| T1 | 23 (57.5) |  |  |
| T2 | 14 (34.1) |  |  |
| T3 | 0 (0) |  |  |
| T4 | 0 (0) |  |  |
| Missing | 3 (7.5) |  |  |
| **Lymph Nodes metastasis n (%)** |  |  |  |
| 0 | 24 (60.0) |  |  |
| 1-3 | 12 (30.0) |  |  |
| ≥4 | 0 (0) | | |
| Missing | 4 (10) |  |  |
| **Histological grading n (%)** |  |  |  |
| I | 2 (5) |  |  |
| II | 27 (67.5) |  |  |
| III | 10 (25) |  |  |
| Missing | 1 (2.5) |  |  |
| **Immunohistochemical status n (%)** |  |  |  |
| ER positive | 40 (100) |  |  |
| PR positive | 40 (100) |  |  |
| HER2 negative | 40 (100) |  |  |
| **Molecular subtype** |  |  |  |
| Luminal A | 25 (62.5) |  |  |
| Luminal B | 13 (32.5) |  |  |
| Missing | 2 (5) |  |  |

<sup>(a)</sup> Numbers may not add up to the total number of patients because of missing values for some variables.
**BC:** breast cancer subjects; **BF:** breast fibroadenoma subjects; **ER:** estrogen receptor; **HD:** healthy donors; **HER2:** human epidermal growth factor receptor 2; **n/a:** not applicable; **PR:** progesterone receptor.

## Notes

### Competing Interest Statement

The authors have declared no competing interest.

## References

1. F. Shan, A. Somasundaram, T. C. Bruno, C. J. Workman, D. A. A. Vignali, Therapeutic targeting of regulatory T cells in cancer. Trends Cancer 8, 944–961 (2022).

2. Y. Togashi, K. Shitara, H. Nishikawa, Regulatory T cells in cancer immunosuppression - implications for anticancer therapy. Nat Rev Clin Oncol 16, 356–371 (2019).

3. G. P. Dunn, A. T. Bruce, H. Ikeda, L. J. Old, R. D. Schreiber, Cancer immunoediting: from immunosurveillance to tumor escape. Nat Immunol 3, 991–998 (2002).

4. J. S. O’Donnell, M. W. L. Teng, M. J. Smyth, Cancer immunoediting and resistance to T cell-based immunotherapy. Nat Rev Clin Oncol 16, 151–167 (2019).

5. J. Blanco-Heredia et al., Converging and evolving immuno-genomic routes toward immune escape in breast cancer. Nat Commun 15, 1302 (2024).

6. N. McGranahan et al., Allele-Specific HLA Loss and Immune Escape in Lung Cancer Evolution. Cell 171, 1259–1271 e1211 (2017).

7. S. Adams et al., Pembrolizumab monotherapy for previously treated metastatic triple-negative breast cancer: cohort A of the phase II KEYNOTE-086 study. Ann Oncol 30, 397–404 (2019).

8. M. E. Gatti-Mays et al., If we build it they will come: targeting the immune response to breast cancer. NPJ Breast Cancer 5, 37 (2019).

9. R. R. Gomis, S. Gawrzak, Tumor cell dormancy. Mol Oncol 11, 62–78 (2017).

10. T. G. Karrison, D. J. Ferguson, P. Meier, Dormancy of mammary carcinoma after mastectomy. J Natl Cancer Inst 91, 80–85 (1999).

11. R. Demicheli et al., Recurrence and mortality dynamics for breast cancer patients undergoing mastectomy according to estrogen receptor status: different mortality but similar recurrence. Cancer Sci 101, 826–830 (2010).

12. M. Colleoni et al., Annual Hazard Rates of Recurrence for Breast Cancer During 24 Years of Follow-Up: Results From the International Breast Cancer Study Group Trials I to V. J Clin Oncol 34, 927–935 (2016).

13. P. F. Denoix, [Nomenclature and classification of cancers based on an atlas]. Acta Unio Int Contra Cancrum 9, 769–771 (1953).

14. A. F. Vieira, F. Schmitt, An Update on Breast Cancer Multigene Prognostic Tests-Emergent Clinical Biomarkers. Front Med (Lausanne) 5, 248 (2018).

15. R. N. Pedersen et al., The Incidence of Breast Cancer Recurrence 10-32 Years After Primary Diagnosis. J Natl Cancer Inst 114, 391–399 (2022).

16. H. Pan et al., 20-Year Risks of Breast-Cancer Recurrence after Stopping Endocrine Therapy at 5 Years. N Engl J Med 377, 1836–1846 (2017).

17. R. L. Siegel, K. D. Miller, A. Jemal, Cancer statistics, 2020. CA Cancer J Clin 70, 7–30 (2020).

18. G. L. Beatty, W. L. Gladney, Immune escape mechanisms as a guide for cancer immunotherapy. Clin Cancer Res 21, 687–692 (2015).

19. M. De Simone et al., Transcriptional Landscape of Human Tissue Lymphocytes Unveils Uniqueness of Tumor-Infiltrating T Regulatory Cells. Immunity 45, 1135–1147 (2016).

20. B. Shang, Y. Liu, S. J. Jiang, Y. Liu, Prognostic value of tumor-infiltrating FoxP3+ regulatory T cells in cancers: a systematic review and meta-analysis. Sci Rep 5, 15179 (2015).

21. A. Tanaka, S. Sakaguchi, Regulatory T cells in cancer immunotherapy. Cell Res 27, 109–118 (2017).

22. R. Saleh, E. Elkord, FoxP3(+) T regulatory cells in cancer: Prognostic biomarkers and therapeutic targets. Cancer Lett 490, 174–185 (2020).

23. S. Liu et al., Prognostic significance of FOXP3+ tumor-infiltrating lymphocytes in breast cancer depends on estrogen receptor and human epidermal growth factor receptor-2 expression status and concurrent cytotoxic T-cell infiltration. Breast Cancer Res 16, 432 (2014).

24. S. A. Perez et al., CD4+CD25+ regulatory T-cell frequency in HER-2/neu (HER)-positive and HER-negative advanced-stage breast cancer patients. Clin Cancer Res 13, 2714–2721 (2007).

25. R. J. deLeeuw, S. E. Kost, J. A. Kakal, B. H. Nelson, The prognostic value of FoxP3+ tumor-infiltrating lymphocytes in cancer: a critical review of the literature. Clin Cancer Res 18, 3022–3029 (2012).

26. J. Stenstrom, I. Hedenfalk, C. Hagerling, Regulatory T lymphocyte infiltration in metastatic breast cancer-an independent prognostic factor that changes with tumor progression. Breast Cancer Res 23, 27 (2021).

27. K. Kos et al., Tumor-educated T(regs) drive organ-specific metastasis in breast cancer by impairing NK cells in the lymph node niche. Cell Rep 38, 110447 (2022).

28. N. R. West et al., Tumour-infiltrating FOXP3(+) lymphocytes are associated with cytotoxic immune responses and good clinical outcome in oestrogen receptor-negative breast cancer. Br J Cancer 108, 155–162 (2013).

29. G. Plitas et al., Regulatory T Cells Exhibit Distinct Features in Human Breast Cancer. Immunity 45, 1122–1134 (2016).

30. J. G. Tate et al., COSMIC: the Catalogue Of Somatic Mutations In Cancer. Nucleic Acids Res 47, D941–D947 (2019).

31. H. Nakagawa et al., Instability of Helios-deficient Tregs is associated with conversion to a T-effector phenotype and enhanced antitumor immunity. Proc Natl Acad Sci U S A 113, 6248–6253 (2016).

32. M. J. Watson et al., Metabolic support of tumour-infiltrating regulatory T cells by lactic acid. Nature 591, 645–651 (2021).

33. R. K. W. Mailer, Alternative Splicing of FOXP3-Virtue and Vice. Front Immunol 9, 530 (2018).

34. N. Goda et al., The ratio of CD8 + lymphocytes to tumor-infiltrating suppressive FOXP3 + effector regulatory T cells is associated with treatment response in invasive breast cancer. Discov Oncol 13, 27 (2022).

35. V. De Rosa et al., Glycolysis controls the induction of human regulatory T cells by modulating the expression of FOXP3 exon 2 splicing variants. Nat Immunol 16, 1174–1184 (2015).

36. S. Junius, et al., Unstable regulatory T cells, enriched for naive and Nrp1(neg) cells, are purged after fate challenge. Sci Immunol 6, (2021).

37. J. Du, et al., FOXP3 exon 2 controls T(reg) stability and autoimmunity. Sci Immunol 7, eabo5407 (2022).

38. L. Demir et al., Predictive and prognostic factors in locally advanced breast cancer: effect of intratumoral FOXP3+ Tregs. Clin Exp Metastasis 30, 1047–1062 (2013).

39. S. Garaud et al., A simple and rapid protocol to non-enzymatically dissociate fresh human tissues for the analysis of infiltrating lymphocytes. J Vis Exp, (2014).

40. W. Sun et al., TSVdb: a web-tool for TCGA splicing variants analysis. BMC Genomics 19, 405 (2018).

41. D. O. Croci et al., Dynamic cross-talk between tumor and immune cells in orchestrating the immunosuppressive network at the tumor microenvironment. Cancer Immunol Immunother 56, 1687–1700 (2007).

42. T. L. Whiteside, The tumor microenvironment and its role in promoting tumor growth. Oncogene 27, 5904–5912 (2008).

43. M. Kanehisa, S. Goto, KEGG: kyoto encyclopedia of genes and genomes. Nucleic Acids Res 28, 27–30 (2000).

44. P. Jayachandran et al., Breast cancer and neurotransmitters: emerging insights on mechanisms and therapeutic directions. Oncogene 42, 627–637 (2023).

45. I. Bozic et al., Accumulation of driver and passenger mutations during tumor progression. Proc Natl Acad Sci U S A 107, 18545–18550 (2010).

46. E. Lee et al., Metabolic stress induces a Wnt-dependent cancer stem cell-like state transition. Cell Death Dis 6, e1805 (2015).

47. K. Takahashi et al., The pluripotent stem cell-specific transcript ESRG is dispensable for human pluripotency. PLoS Genet 17, e1009587 (2021).

48. K. Maskalenka et al., NANOGP1, a tandem duplicate of NANOG, exhibits partial functional conservation in human naive pluripotent stem cells. Development 150, (2023).

49. T. Wu et al., clusterProfiler 4.0: A universal enrichment tool for interpreting omics data. Innovation (Camb) 2, 100141 (2021).

50. L. Beumers et al., Clonal heterogeneity in ER+ breast cancer reveals the proteasome and PKC as potential therapeutic targets. NPJ Breast Cancer 9, 97 (2023).

51. C. Yoon et al., PI3K/Akt pathway and Nanog maintain cancer stem cells in sarcomas. Oncogenesis 10, 12 (2021).

52. M. Karami Fath et al., PI3K/Akt/mTOR signaling pathway in cancer stem cells. Pathol Res Pract 237, 154010 (2022).

53. N. McGranahan et al., Clonal neoantigens elicit T cell immunoreactivity and sensitivity to immune checkpoint blockade. Science 351, 1463–1469 (2016).

54. C. Valero et al., The association between tumor mutational burden and prognosis is dependent on treatment context. Nat Genet 53, 11–15 (2021).

55. A. Walens et al., Adaptation and selection shape clonal evolution of tumors during residual disease and recurrence. Nat Commun 11, 5017 (2020).

56. M. S. Lawrence et al., Discovery and saturation analysis of cancer genes across 21 tumour types. Nature 505, 495–501 (2014).

57. S. Nik-Zainal et al., Landscape of somatic mutations in 560 breast cancer whole-genome sequences. Nature 534, 47–54 (2016).

58. S. Nik-Zainal et al., The life history of 21 breast cancers. Cell 149, 994–1007 (2012).

59. M. B. Burns et al., APOBEC3B is an enzymatic source of mutation in breast cancer. Nature 494, 366–370 (2013).

60. L. B. Alexandrov et al., Signatures of mutational processes in human cancer. Nature 500, 415–421 (2013).

61. S. Haricharan et al., Loss of MutL Disrupts CHK2-Dependent Cell-Cycle Control through CDK4/6 to Promote Intrinsic Endocrine Therapy Resistance in Primary Breast Cancer. Cancer Discov 7, 1168–1183 (2017).

62. M. Anurag et al., Comprehensive Profiling of DNA Repair Defects in Breast Cancer Identifies a Novel Class of Endocrine Therapy Resistance Drivers. Clin Cancer Res 24, 4887–4899 (2018).

63. X. Wang et al., TIMEDB: tumor immune micro-environment cell composition database with automatic analysis and interactive visualization. Nucleic Acids Res 51, D1417–D1424 (2023).

64. T. C. Brown, J. Jiricny, Repair of base-base mismatches in simian and human cells. Genome 31, 578–583 (1989).

65. M. Haruna et al., The impact of CCR8+ regulatory T cells on cytotoxic T cell function in human lung cancer. Sci Rep 12, 5377 (2022).

66. T. Wang et al., CCR8 blockade primes anti-tumor immunity through intratumoral regulatory T cells destabilization in muscle-invasive bladder cancer. Cancer Immunol Immunother 69, 1855–1867 (2020).

67. H. Kennecke et al., Metastatic behavior of breast cancer subtypes. J Clin Oncol 28, 3271–3277 (2010).

68. N. A. Soliman, S. M. Yussif, Ki-67 as a prognostic marker according to breast cancer molecular subtype. Cancer Biol Med 13, 496–504 (2016).

69. A. C. Wolff et al., Human Epidermal Growth Factor Receptor 2 Testing in Breast Cancer: American Society of Clinical Oncology/College of American Pathologists Clinical Practice Guideline Focused Update. J Clin Oncol 36, 2105–2122 (2018).

70. A. Raugh, D. Allard, M. Bettini, Nature vs. nurture: FOXP3, genetics, and tissue environment shape Treg function. Front Immunol 13, 911151 (2022).

71. L. Wang et al., PARP1 in Carcinomas and PARP1 Inhibitors as Antineoplastic Drugs. Int J Mol Sci 18, (2017).

72. A. Glaviano et al., PI3K/AKT/mTOR signaling transduction pathway and targeted therapies in cancer. Mol Cancer 22, 138 (2023).

73. P. M. K. Westcott et al., Mismatch repair deficiency is not sufficient to elicit tumor immunogenicity. Nat Genet 55, 1686–1695 (2023).

74. M. Efremova, F. Finotello, D. Rieder, Z. Trajanoski, Neoantigens Generated by Individual Mutations and Their Role in Cancer Immunity and Immunotherapy. Front Immunol 8, 1679 (2017).

75. P. Baldominos et al., Quiescent cancer cells resist T cell attack by forming an immunosuppressive niche. Cell 185, 1694–1708 e1619 (2022).

76. S. Adams et al., A Multicenter Phase II Trial of Ipilimumab and Nivolumab in Unresectable or Metastatic Metaplastic Breast Cancer: Cohort 36 of Dual Anti-CTLA-4 and Anti-PD-1 Blockade in Rare Tumors (DART, SWOG S1609). Clin Cancer Res 28, 271–278 (2022).

77. R. N. Amaria et al., Neoadjuvant immune checkpoint blockade in high-risk resectable melanoma. Nat Med 24, 1649–1654 (2018).

78. M. D. Hellmann et al., Nivolumab plus Ipilimumab in Lung Cancer with a High Tumor Mutational Burden. N Engl J Med 378, 2093–2104 (2018).

79. J. A. Kyte et al., ICON: a randomized phase IIb study evaluating immunogenic chemotherapy combined with ipilimumab and nivolumab in patients with metastatic hormone receptor positive breast cancer. J Transl Med 18, 269 (2020).

80. A. Ribas, J. D. Wolchok, Cancer immunotherapy using checkpoint blockade. Science 359, 1350–1355 (2018).

81. C. W. Elston, I. O. Ellis, Pathological prognostic factors in breast cancer. I. The value of histological grade in breast cancer: experience from a large study with long-term follow-up. Histopathology 19, 403–410 (1991).

82. P. H. Tan et al., The 2019 World Health Organization classification of tumours of the breast. Histopathology 77, 181–185 (2020).

83. W. J. Kent et al., The human genome browser at UCSC. Genome Res 12, 996–1006 (2002).

84. C. Fresno, E. A. Fernandez, RDAVIDWebService: a versatile R interface to DAVID. Bioinformatics 29, 2810–2811 (2013).

85. W. Walter, F. Sanchez-Cabo, M. Ricote, GOplot: an R package for visually combining expression data with functional analysis. Bioinformatics 31, 2912–2914 (2015).

86. G. Yu, L. G. Wang, Y. Han, Q. Y. He, clusterProfiler: an R package for comparing biological themes among gene clusters. OMICS 16, 284–287 (2012).

87. A. Colaprico et al., TCGAbiolinks: an R/Bioconductor package for integrative analysis of TCGA data. Nucleic Acids Res 44, e71 (2016).

88. A. Mayakonda, D. C. Lin, Y. Assenov, C. Plass, H. P. Koeffler, Maftools: efficient and comprehensive analysis of somatic variants in cancer. Genome Res 28, 1747–1756 (2018).

89. M. S. Lawrence et al., Mutational heterogeneity in cancer and the search for new cancer- associated genes. Nature 499, 214–218 (2013).

90. R. Gaujoux, C. Seoighe, A flexible R package for nonnegative matrix factorization. BMC Bioinformatics 11, 367 (2010).

